# SARM1 is responsible for calpain-dependent dendrite degeneration in mouse hippocampal neurons

**DOI:** 10.1101/2022.04.27.489702

**Authors:** Takashi Miyamoto, Chaeyoung Kim, Johann Chow, Jason C Dugas, Jack DeGroot, Alex L Bagdasarian, Arun P Thottumkara, Martin Larhammar, Meredith EK Calvert, Brian M Fox, Joseph W Lewcock, Lesley A Kane

## Abstract

Sterile alpha and TIR motif containing 1 (SARM1) is a critical regulator of axon degeneration that acts through hydrolysis of NAD^+^ following injury. Recent work has defined the mechanisms underlying SARM1’s catalytic activity and advanced our understanding of SARM1 function in axons, yet the role of SARM1 signaling in other compartments of neurons is still not well understood. Here we show in cultured hippocampal neurons that endogenous SARM1 is present in axons, dendrites and cell bodies and that direct activation of SARM1 by the neurotoxin Vacor causes not just axon degeneration, but degeneration of all neuronal compartments. In contrast to the axon degeneration pathway defined in dorsal root ganglia (DRGs), SARM1-dependent hippocampal axon degeneration in vitro is not sensitive to calpain inhibition whereas dendrite degeneration downstream of SARM1 is calpain-dependent in this cell type. This data indicates SARM1 plays a critical role in neurodegeneration outside of axons and elucidates divergent pathways leading to degeneration in hippocampal axons and dendrites.

## Introduction

Distal axon degeneration following injury follows a pattern that was originally termed Wallerian degeneration (1, 2). For many years it was believed that this degeneration was passive, but this changed after the identification of the Wallerian Degeneration Slow (WLDS) mouse strain that displayed significantly delayed axon degeneration following injury (3). The WLDS mutation was subsequently shown to be a fusion of ubiquitination factor E4B (UBE4B) and nicotinamide mononucleotide adenylyltransferase 1 (NMNAT1) (4). This fusion protein persists in damaged axons and has been shown to be protective in injury models of cultured dorsal root ganglion (DRG) or superior cervical ganglion (SCG) neurons and mouse sciatic nerves (5–7). NMNAT1 is an enzyme that converts nicotinamide mononucleotide (NMN) into NAD^+^ that is usually localized to the nucleus, but the UBE4B-NMNAT1 fusion protein is diffusely localized and stabilized. This results in active enzyme that persists in damaged axons and preserves NAD^+^ levels following injury (8). Further studies in SCG neurons revealed that the endogenous protein responsible for maintaining NAD^+^ levels in axons is another NMNAT isoform, NMNAT2, which is rapidly lost upon axon injury (9).

A subsequent forward genetic screen in *Drosophila melanogaster* led to the discovery of the first mutations that slowed Wallerian degeneration (10). This study identified Sterile alpha and Armadillo motif (*dSarm*) and its mouse ortholog Sterile alpha and TIR motif-containing protein 1 (*Sarm1*) as a novel regulator of axon degeneration. Deletion of *dSarm*/*Sarm1* in flies, primary mouse cortical, SCG, DRG neurons, and mouse sciatic nerve delayed axon degeneration following injury. The mechanism by which SARM1 regulates axon degeneration became clearer upon the discovery of a non-canonical NAD^+^ hydrolase function of its TIR domain (11). Isolated recombinant or cellular SARM1 hydrolyzes NAD^+^ into nicotinamide, adenosine diphosphate ribose (ADPR), and cyclic-ADPR (cADPR) (11–14). SARM1’s intrinsic NAD^+^ hydrolase activity, along with the known roles of WLDS and NMNAT2 in NAD^+^ metabolism, solidified SARM1-dependent NAD^+^ hydrolysis as a critical step in the axon degeneration pathway (1, 2). Taken together, this data has led to a model in which degradation of NMNAT2 lowers NAD^+^ and increases NMN following axonal injury (15–17). NMN then directly activates SARM1, leading to a further reduction of NAD^+^ (2, 14, 18–25). Structural investigations confirmed an allosteric binding pocket in SARM1, normally occupied by NAD^+^ but replaced by NMN to induce hydrolase activity (2, 14, 18, 21, 22, 24, 26, 27). It has been hypothesized that other pro-degenerative stimuli may similarly alter the NAD^+^/NMN ratio to favor activation of SARM1 (15, 18, 24, 27).

Research on SARM1 has focused on its role in axon degeneration following axotomy or treatment with chemotherapeutics (1, 2, 28) and therefore our understanding of mammalian SARM1 function comes predominantly from studies in DRG/SCG neurons as a model system (1, 28, 29). This specialized population of sensory neurons have a single axon that bifurcates into two branches, which has limited our understanding of SARM1 function in other neuronal compartments. Wallerian degeneration of dendrites has been reported in *Drosophila* (30, 31), but there is limited evidence for the pathway in dendrites of mammalian neurons. In this study, we outline the degenerative properties of SARM1 in mouse primary hippocampal neurons that have complex morphology and processes. Using these neurons we show the presence of SARM1 protein and activity outside of axons and define a role for SARM1 in dendrite degeneration through activation of calpain proteases.

## Results

### SARM1 is present and functional in all neuronal compartments

To better define the role of SARM1 in CNS neurons, we began by determining the SARM1 localization in cultured primary hippocampal neurons. Immunostaining for endogenous SARM1 in *wildtype* neurons revealed punctate, but specific signal throughout the cells (Figure 1A – Top row) which was absent in neurons from *Sarm1*-knockout (*Sarm1KO*) mice (Figure 1A – Middle row). We next examined a 1:1 ratio mixed culture of *wildtype* and *Sarm1KO* neurons which allowed us to both confirm the specificity of the SARM1 antibody by observing cells with (Figure 1A – Bottom row, arrow) and without (Figure 1A – Bottom row, arrowhead) signal and to better observe morphological details of each SARM1-positive neuron. Using super-resolution microscopy, we then investigated the presence of SARM1 in neuronal compartments marked by axonal (SMI312) and dendrite (MAP2) markers and found that endogenous SARM1 is present in axons, dendrites, and cell bodies (Figure 1B – zoomed in image and line graph).

**Figure 1.**
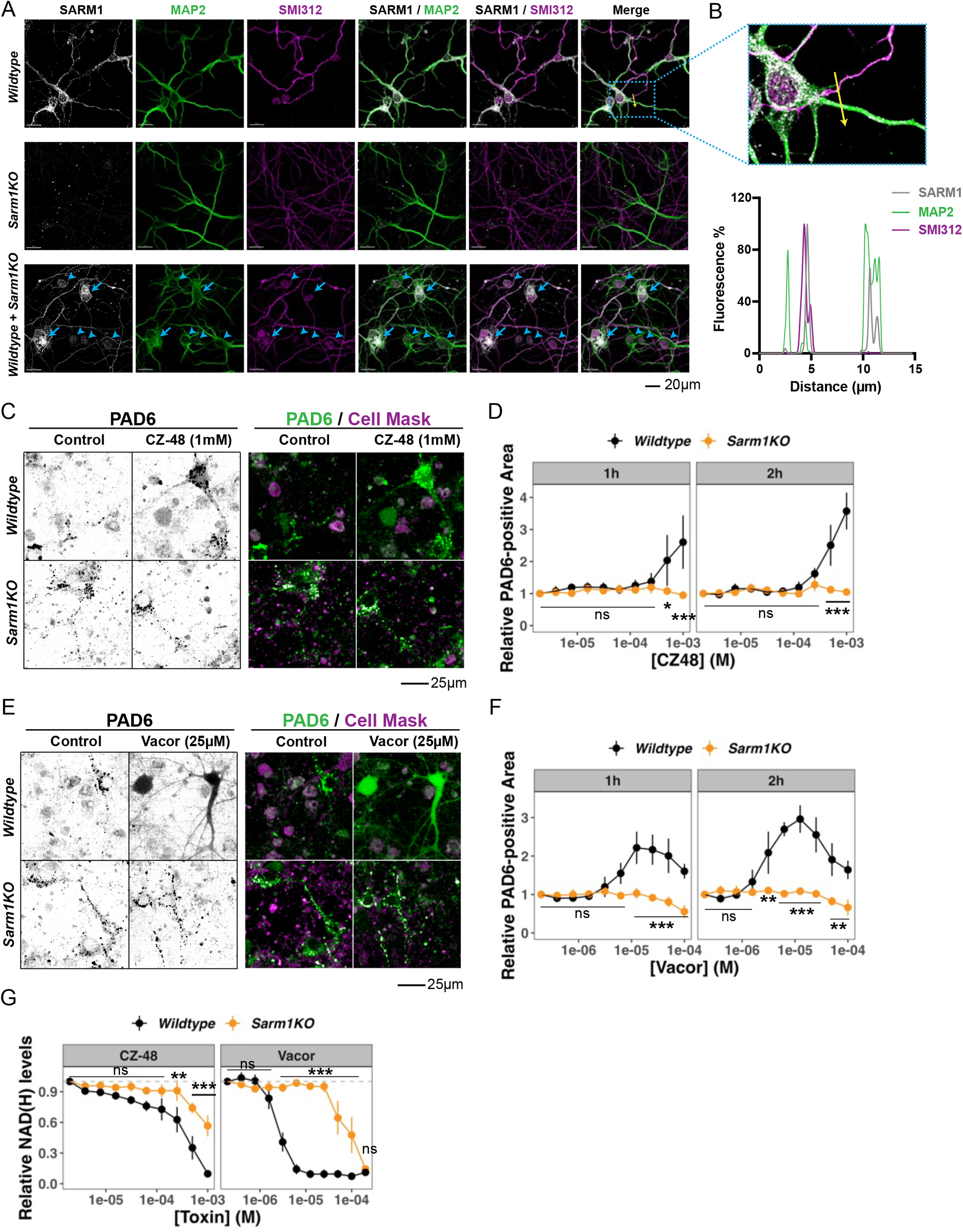
SARM1 is present and can be activated in all neuronal compartments. (**A**) Representative images of the primary cultured hippocampal neurons (“neurons”) from *wildtype* and *Sarm1KO* mice, and the mixed culture of the two genotypes that were immunostained for endogenous SARM1 (grayscale), axonal (SMI312, magenta), and dendritic (MAP2, green) marker proteins. In the mixed neuron culture, blue arrows and arrowheads point at *wildtype* and *Sarm1KO* neurons, respectively, that were identified based on the SARM1 immunostaining. (**B**) A zoomed-in image of the *wildtype* neurons in (**A**) - the area defined by the blue dashed box in the merged image of *wildtype* neurons (top), and the line graph that indicates the relative intensity of SARM1, MAP2, and SMI312 immunostaining along the yellow arrow indicated in the zoomed-in image (bottom). (**C-F**) SARM1 activity in *wildtype* and *Sarm1KO* neurons treated with CZ-48 (**C** and **D**) or Vacor (**E** and **F**) shown by both representative images at 1 hour (**C** and **E,** grayscale (left) and overlaid with CellMask staining (right)) and quantification of PAD6-positive area for 1 or 2 hours (**D** and **F**). (**G**) Cellular NAD(H) levels of *wildtype* and *Sarm1KO* neurons that were treated with increasing concentrations of CZ-48 or Vacor for 16 hours. Statistical significance of differences among groups was determined by two-way ANOVA using genotype and toxin doses as two independent variables. When the effect of genotype was significant (P < 0.05), Tukey’s post hoc test was performed and P values that are less than 0.05 for comparisons between the two genotypes at the specific toxin doses are shown. Numbers of biologically independent experiments (n) are (**D, F**) 3 and (**G**) 3 (CZ-48) and 6 (Vacor). Data represent the mean ± SEM, * P < 0.05, ** P < 0.01, *** P < 0.001, ns P ≥ 0.05.

We next sought to test SARM1 activity in each neuronal compartment following direct activation by CZ-48 or Vacor, two cell permeable compounds that have been reported to mimic NMN, an endogenous SARM1 activator (20, 23, 32). CZ-48 is an inhibitor of CD38, another NADase that was unexpectedly found to activate SARM1 (20), and Vacor was used as a rodenticide and its accidental ingestion can cause peripheral and central neuropathy and/or diabetes mellitus in humans (33, 34). While CZ-48 is an NMN analog, Vacor requires processing by nicotinamide phosphoribosyltransferase (NAMPT) to produce the NMN analog VMN, which directly activates SARM1 (20, 23, 32). To directly observe SARM1 activity, we employed PC6/PAD6, a live-cell SARM1 activity probe. PAD6 is formed as an adduct of PC6 and ADPR generated by SARM1 upon NAD^+^ hydrolysis that causes a red shift in fluorescence and allows for the direct measurement of SARM1 activity in live cells (35). Neurons were pre-loaded with PC6, treated with a wide concentration range of SARM1 activators, and then imaged for PAD6 signal at two time points – 1, and 2 hours after the addition of SARM1 agonists. Following SARM1 activation with CZ-48 (Figure 1C-D) or Vacor (Figure 1E-F), *wildtype*, but not *Sarm1KO*, neurons displayed diffuse fluorescent signal throughout cell bodies and neurites, indicating the signal was induced by SARM1-dependent PAD6 formation across all the neuronal compartments. The PAD6 signal was also ablated by co-treatment with a SARM1 inhibitor (36) in *wildtype* neurons (Supplemental Figure 1-1A-B). PC6-loaded *wildtype* and *Sarm1KO* neurons both displayed some punctate background signal in the control conditions that overlapped with lysosomes (Supplemental Figure 1-1C), suggesting protonation of PC6 in acidic lysosomes that was independent of SARM1 activity. However, this background signal could be readily separated from the PAD6 signal observed throughout neurons (Figure 1C, E). Following activation of SARM1 in hippocampal neurons by either CZ-48 or Vacor, total cellular NAD(H) was lost in a SARM1-dependent manner (Figure 1G). This ubiquitous PAD6 signal and complete loss of cellular NAD(H) indicates that SARM1 is functional and can be activated both inside and outside of axons in hippocampal neurons.

### Direct SARM1 activation results in degeneration of multiple neuronal compartments

To gain a better understanding of the consequences of SARM1 activation in distinct subcellular compartments, we directly activated SARM1 using a wide concentration range of CZ-48 or Vacor and then assessed the extent of degeneration at a single timepoint (16 hours) by immunostaining for markers of axons (SMI312) and dendrites (MAP2), as well as quantifying nuclear condensation (DAPI). Both compounds resulted in a dose-dependent degeneration of both axons and dendrites in *wildtype*, but not *Sarm1KO*, neurons (Figure 2A-D). Neurons treated with very high (≥ 50µM) concentrations of Vacor also showed SARM1-independent degeneration, presumably a result of off-target toxicity (Figure 2D). Degeneration of neuronal processes following CZ-48/Vacor treatment was similarly attenuated by pharmacological inhibition of SARM1 activity (Supplemental Figure 2-1A). Vacor and CZ-48 also caused a SARM1-dependent increase in condensed nuclei, suggesting that in this setting SARM1 activation can be sufficient to drive cell death in addition to neurite degeneration (Figure 2A-D).

**Figure 2.**
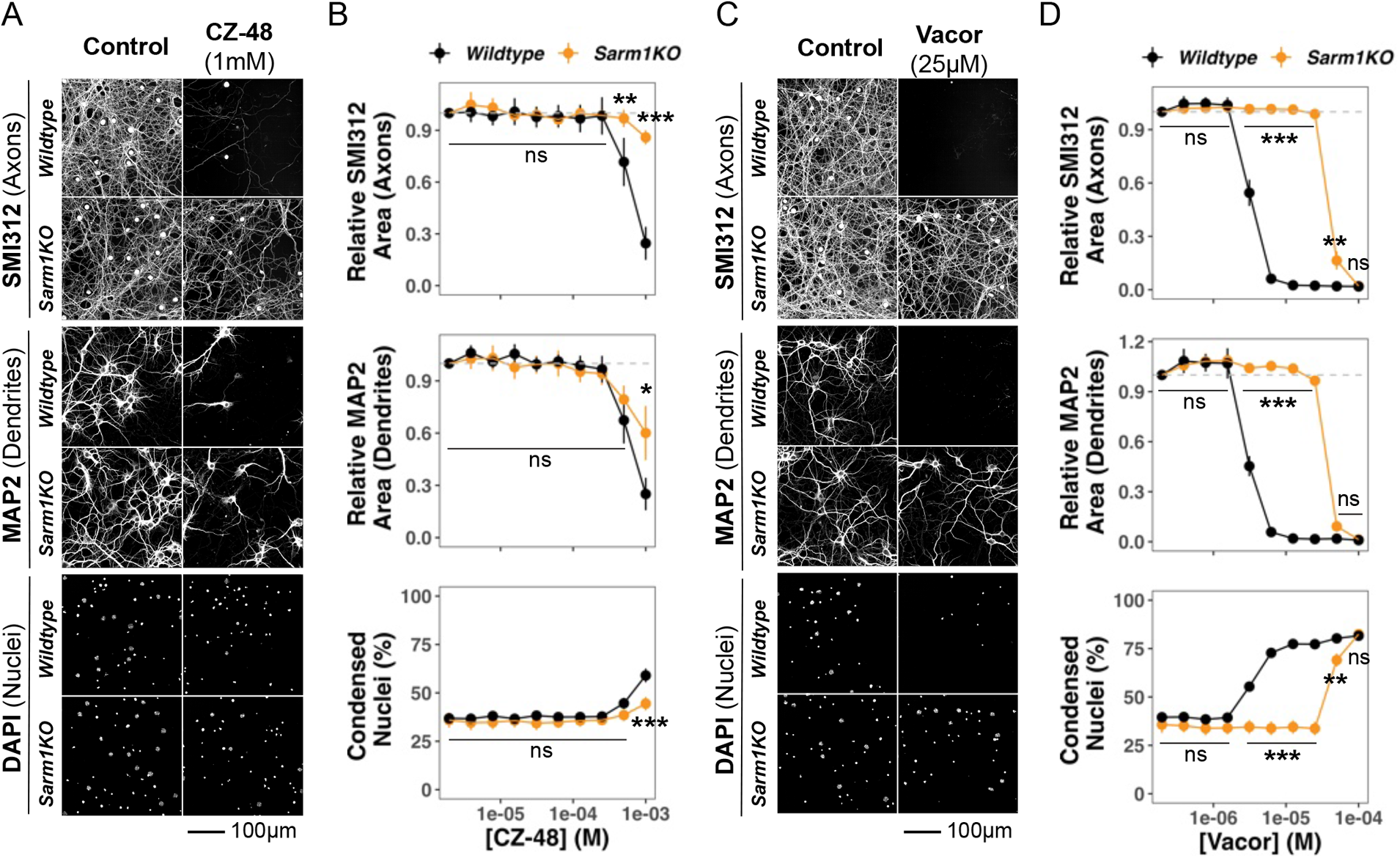
SARM1 activation degenerates all neuronal compartments. Representative images and relative levels of immunostained axons and dendrites, and proportion of condensed nuclei in *wildtype* and *Sarm1KO* neurons that were treated with increasing concentrations of (**A, B**) CZ-48 or (**C, D**) Vacor for 16 hours. All representative images from the same group were taken in the same field of the same neuronal culture. Statistical significance of differences among groups was determined by two-way ANOVA using genotype and toxin doses as two independent variables. When the effect of genotype was significant (P < 0.05), Tukey’s post hoc test was performed and P values that are less than 0.05 for comparisons between the two genotypes at the specific toxin doses are shown. Numbers of biologically independent experiments (n) are (**B**) 5, and (**D**) 4. Data represent the mean ± SEM, * P < 0.05, ** P < 0.01, *** P < 0.001, ns P ≥ 0.05.

All neuronal compartments (axons, dendrites, and cell bodies) had the same relative sensitivity to CZ-48 and Vacor, indicating that no one compartment is more sensitive to SARM1 activation. Vacor was a more potent activator with maximal degeneration occurring at 6.25 µM, while CZ-48 did not reach maximal levels even at 1 mM (Figure 2B and D). However, simultaneous addition of the NAMPT inhibitor FK866 potentiated the degeneration of all neuronal compartments observed following CZ-48 treatment (Supplemental Figure 2-1B), presumably a result of more robust lowering of NAD^+^ levels (14, 18, 22). The addition of FK866 sensitized axons, dendrites and cell bodies to CZ-48 to the same degree with degeneration evident at concentrations ≥ 0.25 mM and maximal degeneration occurring at 0.5 mM (Supplemental Figure 2-1B).

### SARM1-mediated dendrite degeneration is calpain-dependent

Calcium (Ca^2+^) influx and activity of calpains (Ca^2+^-dependent proteases), are known contributors to axon degeneration following injury or SARM1 activation in DRG/SCG neurons (37–42). Based on these findings, we investigated the involvement of calpains in SARM1-dependent degeneration of hippocampal neuron compartments by co-treating neurons with Vacor and several commonly used calpain inhibitors (Supplemental Figure 3-1 A-B). Calpain inhibitor III, Calpeptin, and PD150606 are relatively selective to calpain-1 and -2, compared to EST and Leupeptin that also inhibit other types of proteases (43–45). Surprisingly, none of the calpain inhibitors prevented Vacor-dependent degeneration of SMI312-positive axons even when used at high concentrations (30 µM) (Figure 3A-B, Supplemental Figure 3-1 A-B). In contrast, Calpain inhibitor III and Calpeptin robustly prevented Vacor-dependent degeneration of MAP2-positive dendrites (Figure 3A-B, Supplemental Figure 3-1 A-B). Calpain inhibition also protected dendrites against direct activation of SARM1 by CZ-48/FK866, further supporting the notion that calpains act downstream of SARM1 (Supplemental Figure 3-1 C-D).

**Figure 3.**
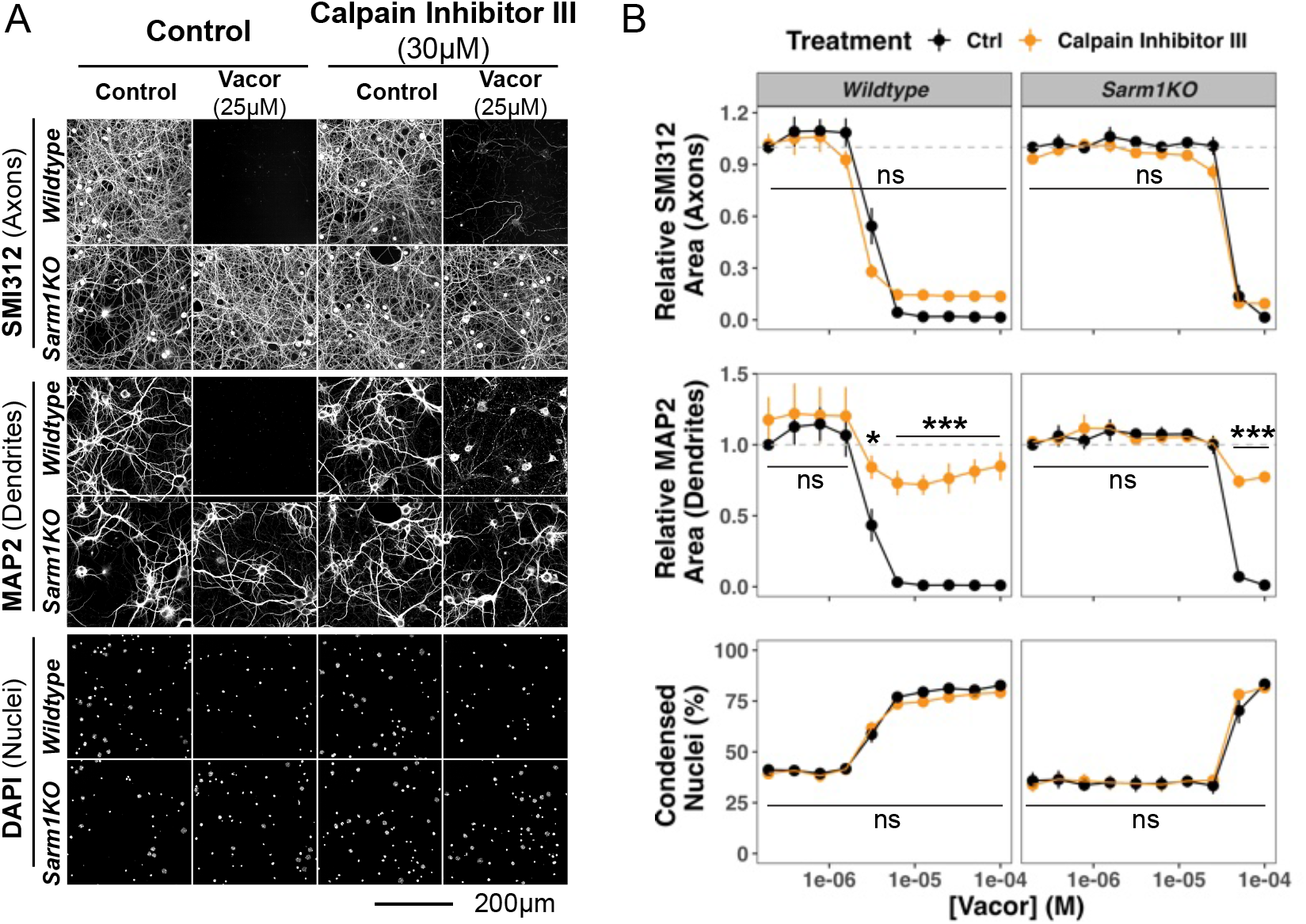
SARM1-induced dendrite degeneration depends on calpains. (**A**) Representative images and (**B**) relative levels of immunostained axons and dendrites, and proportion of condensed nuclei in *wildtype* and *Sarm1KO* neurons that were treated with various concentrations of Vacor for 16 hours. Some neurons were co-treated with calpain inhibitor III (30µM). Statistical significance of effects by pharmacological treatment was determined by two-way ANOVA using Calpain Inhibitor III treatment and Vacor doses as two independent variables. When the effect of the inhibitor treatment was significant (P < 0.05), Tukey’s post hoc test was performed and P values that are less than 0.05 for the effect of inhibitor treatment compared to the control group at the specific toxin doses are shown in figure panels. Numbers of biologically independent experiments (n) are 4. Data represent the mean ± SEM, * P < 0.05, ** P < 0.01, *** P < 0.001, ns P ≥ 0.05.

To determine whether calpain activation and dendrite degeneration is dependent on caspases as was observed in DRG axons following lesion or trophic factor withdrawal (46, 47), we next tested the pan-caspase inhibitor Q-VD-OPh. Caspase inhibition had no effect on SARM1-dependent dendrite degeneration, demonstrating specificity of calpain proteases in this process and suggesting distinct upstream signals lead to calpain activation in this context (Supplemental Figure 3-2). Interestingly, Q-VD-OPh also failed to protect axons of hippocampal neurons from SARM1-dependent degeneration (Supplemental Figure 3-2).

To confirm that calpain activation occurs downstream of SARM1, we delayed the addition of either a SARM1 or calpain inhibitor until after activating SARM1 (See Figure 4A for experimental design). Based on the robust protection of dendrites (Figure 3 and Supplemental Figure 3-1) and known selectivity and potency, Calpain inhibitor III was chosen for subsequent experiments. Delayed addition of a SARM1 inhibitor protected dendrites when added within two hours of Vacor treatment (Figure 4B and Figure 4C, orange shading), whereas Calpain inhibitor III protected dendrites if added up to three hours after Vacor treatment (Figure 4B and Figure 4C, blue shading). The window of one hour where calpain inhibition was still protective beyond SARM1 inhibition suggests that calpains are activated downstream of SARM1 and, once active, are not reversable by SARM1 inhibition. Examination of SARM1 activity following treatment with a calpain inhibitor provided additional evidence that SARM1 acts upstream of calpain activation, as PAD6 fluorescence following co-treatment of hippocampal neurons with CZ-48 or Vacor and Calpain inhibitor III was not significantly different than treatment with SARM1 activators alone (Figure 4D-E).

**Figure 4.**
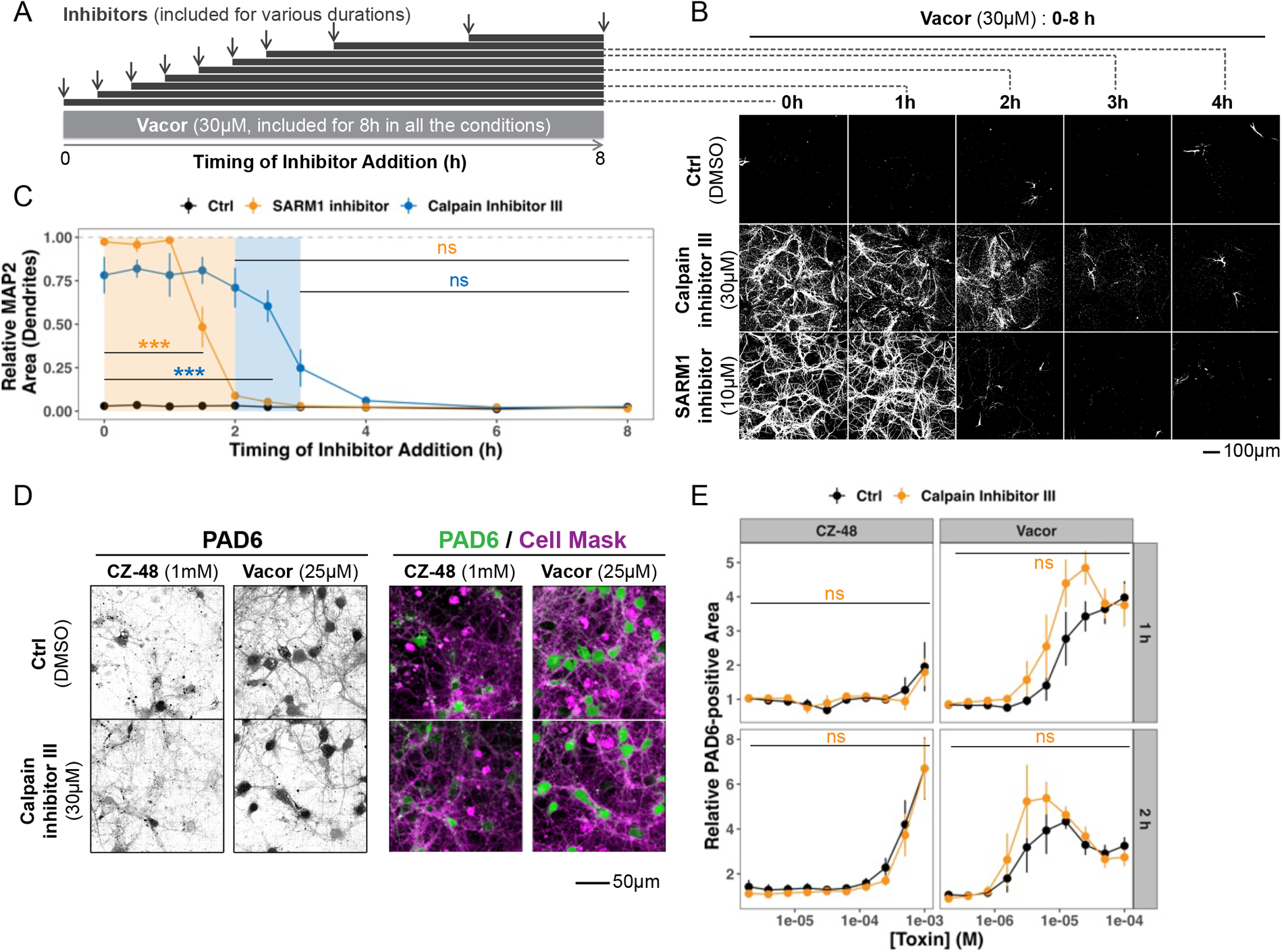
Calpain activation is downstream of SARM1. (**A**) Schematic diagram of experiment in which inhibitors were applied at multiple time points after beginning Vacor treatment. (**B**) Representative images and (**C**) relative levels of immunostained dendrites in *wildtype* neurons that were treated with Vacor for 8 hours with either Calpain Inhibitor III or SARM1 inhibitor added at increasing durations after Vacor treatment as indicated in the panel (**A**). (**D**) Representative images at 1 hour and (**E**) relative PAD6 fluorescence levels of PC6-loaded neurons that were treated with increasing concentrations of CZ-48 or Vacor for 1 or 2 hours with or without Calpain Inhibitor III (30µM). Statistical significance of effects by inhibitors was determined by two-way ANOVA using (**C**) inhibitor treatment and the timing of inhibitor addition, or (**E**) Calpain Inhibitor III treatment and Vacor doses as two independent variables. When the effect of the inhibitor treatment was significant (P < 0.05), Tukey’s post hoc test was performed and P values that are less than 0.05 for the effect of inhibitor treatment at the specific toxin doses are shown in a color-matched manner in figure panels. Numbers of biologically independent experiments (n) are (**C**) 4 or (**E**) 5. Data represent the mean ± SEM, * P < 0.05, ** P < 0.01, *** P < 0.001, ns P ≥ 0.05.

Multiple waves of calcium influx have been described in axon lesion studies, a first transient wave that occurs rapidly after injury and a second prolonged one that occurs hours after injury and is immediately followed by axon fragmentation (39, 40). Given the requirement for calcium influx in calpain activation, we hypothesized that SARM1 activity contributes to calcium influx in hippocampal neurons. To test this, we monitored intracellular calcium levels by Fluo4 intensity following Vacor treatment. Vacor had no effect on calcium influx at early time points (≤ 1h) but resulted in a significant increase in Fluo4 signal in *wildtype* neurons at later time points (≥ 2h) (Figure 5A-B and see Supplemental Figure 5-1A for all timepoints and treatments). Interestingly, Fluo4 signal did not increase at any time point in either *Sarm1KO* neurons or *wildtype* neurons treated with either EGTA or a SARM1 inhibitor, indicating that it is both extracellular calcium and SARM1 activity-dependent (Figure 5A-B and see Supplemental Figure 5-1A for all timepoints and treatments). We next sought to determine the calcium channels acting downstream of SARM1 activation. Among several voltage-gated calcium channels (Cav) blockers, Cilnidipine prevented a large portion of calcium influx induced by Vacor (Figure 5C-D), whereas the other blockers, including Nifedipine, had no effect on calcium influx (Supplemental Figure 5-1B). This suggests that Cav2 channels conduct calcium influx induced by SARM1 activity in neurons as Cilnidipine and Nifedipine block Cav1/2 and Cav1 channels respectively (48).

**Figure 5.**
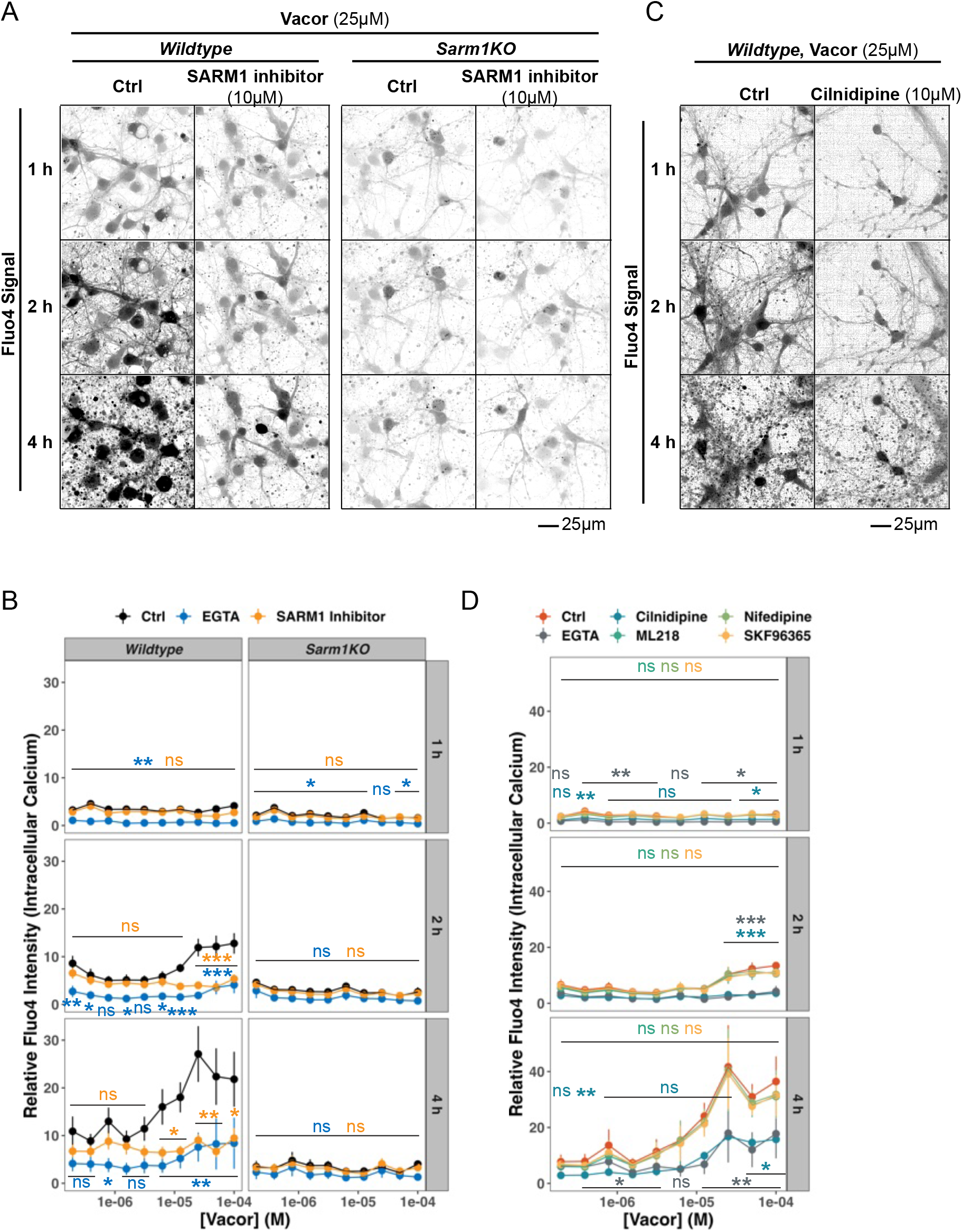
SARM1 activation increases intracellular calcium. (**A**) Representative images of Fluo4 intensity (intracellular calcium) in *wildtype* or *Sarm1KO* neurons treated with 25µM of Vacor at 1, 2, and 4 hours with or without SARM1 inhibitor (10µM). (**B**) Quantification of relative Fluo4 intensity at 1, 2 and 4 hours following treatment with increasing concentrations of Vacor with or without SARM1 inhibitor (10µM) or EGTA (3 mM, See Supplemental Figure 5-1A for EGTA images). (**C**) Representative images of Fluo4 intensity in *wildtype* neurons with or without Cilnidipine (10µM). (**D**) Quantification of relative Fluo4 intensity at 1, 2 and 4 hours following treatment with increasing concentrations of Vacor with or without the indicated calcium channel blockers (See Supplemental Figure 5-1B for other representative images). Statistical significance of effects by inhibitors was determined by two-way ANOVA using inhibitor treatment (including Calpain Inhibitor III, EGTA, SARM1 inhibitor, or Cilnidipine treatment and others) and Vacor doses as two independent variables. When the effect of the inhibitor treatment was significant (P < 0.05), either (**B**) Tukey’s or (**D**) Dunnett’s post hoc test was performed and P values that are less than 0.05 for the effect of inhibitor treatment at the specific Vacor doses are shown in a color-matched manner in figure panels. Numbers of biologically independent experiments (n) are (**B**) 4 and (**D**) 3. Data represent the mean ± SEM, * P < 0.05, ** P < 0.01, *** P < 0.001, ns P ≥ 0.05.

### Calpains are downstream executors of SARM1 degeneration in dendrites, but not axons

Several axon-specific Calpain substrates have been identified including neurofilament light (Nf-L) in axon degeneration in DRG neurons (37, 41, 49). Consistent with previous reports, we observed Nf-L cleavage upon SARM1 activation by Vacor treatment in cultured mouse DRG neurons, which was prevented following addition of Calpain inhibitor III and EGTA (Figure 6A-B). We next tested whether similar Nf-L processing occurred in hippocampal neurons. Surprisingly, despite robust axon degeneration following Vacor treatment in hippocampal neurons (Figure 2), no calpain-dependent processing of Nf-L occurred (Figure 6C-D). Robust and specific Nf-L expression in axons of these neurons was confirmed by immunostaining, indicating that this discrepancy was not a result of changes in subcellular localization (Supplemental Figure 6-1A). This lack of Nf-L processing by calpains is consistent with the lack of axonal protection afforded by calpain inhibition in this paradigm.

**Figure 6.**
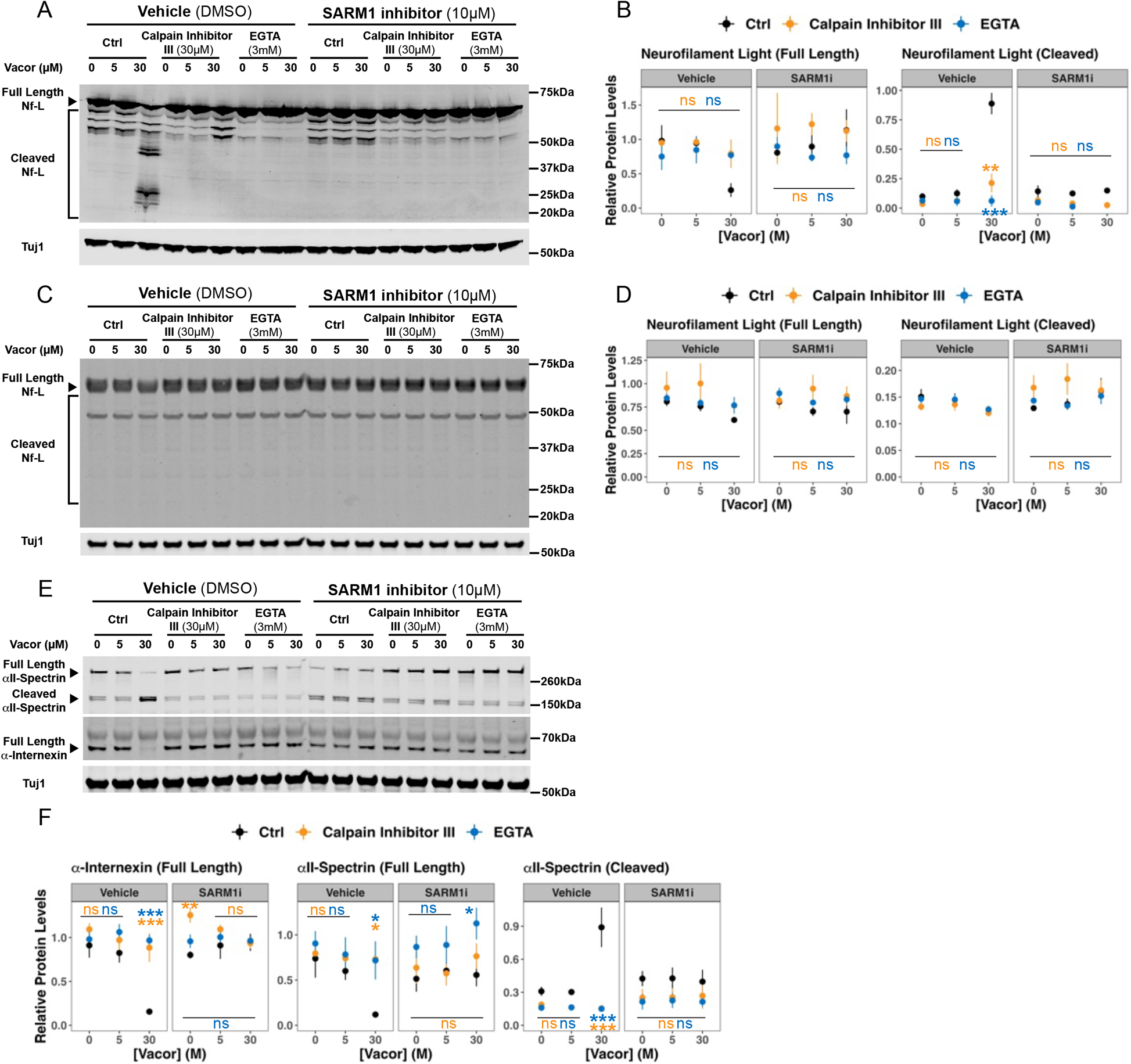
Calpains are downstream executors of SARM1 degeneration in dendrites, but not axons. (**A**) Representative western blot images and (**B**) quantification of relative levels of full-length and cleaved neurofilament light (Nf-L) in mouse DRG neurons treated with increasing concentrations of Vacor with or without SARM1 inhibitor (10µM), Calpain inhibitor III (30µM), or EGTA (3mM). (**C**) Representative western blot images and (**D**) quantification of relative levels of full-length and cleaved neurofilament light (Nf-L) in mouse hippocampal neurons treated with increasing concentrations of Vacor with or without SARM1 inhibitor (10µM), Calpain inhibitor III (30µM) or EGTA (3mM). (**E**) Representative western blot images and (**F**) relative levels of full length α-Internexin, full length, and cleaved αII-Spectrin under the same conditions as (**C-D**). Statistical significance of effects by inhibitors were determined by two-way ANOVA using inhibitor and Vacor treatment as two independent variables. When the effect of the inhibitor treatment was significant (P < 0.05), Tukey’s post hoc test was performed and P values that are less than 0.05 for the effect of pharmacological treatment compared to the control group at the specific Vacor doses are shown in a color-matched manner in figure panels. Numbers of biologically independent experiments (n) are (**B**) 3, and (**D, F**) 4. **-** Data represent the mean ± SEM, * P < 0.05, ** P < 0.01, *** P < 0.001, ns P ≥ 0.05.

As dendrites were specifically protected by calpain inhibition following Vacor we moved on to measure the cleavage of αII-spectrin and α-internexin, two calpain substrates that are present in both dendrites and axons of hippocampal neurons (50, 51) (Supplemental Figure 6-1B-C). Both αII-spectrin and α-internexin were substantially degraded by treatment with 30 µM Vacor for 3 hours (Figure 6E-F). This processing was completely blocked by co-treatment with a SARM1 or calpain inhibitor, confirming that calpain proteolytic activity occurs following Vacor treatment (Figure 6E-F). Since the processing is specific to proteins found in dendrites, calpains appear to only be activated downstream of SARM1 in dendrites and not in axons where an alternative degeneration pathway may exist downstream of SARM1 (see model in Figure 7).

**Figure 7.**
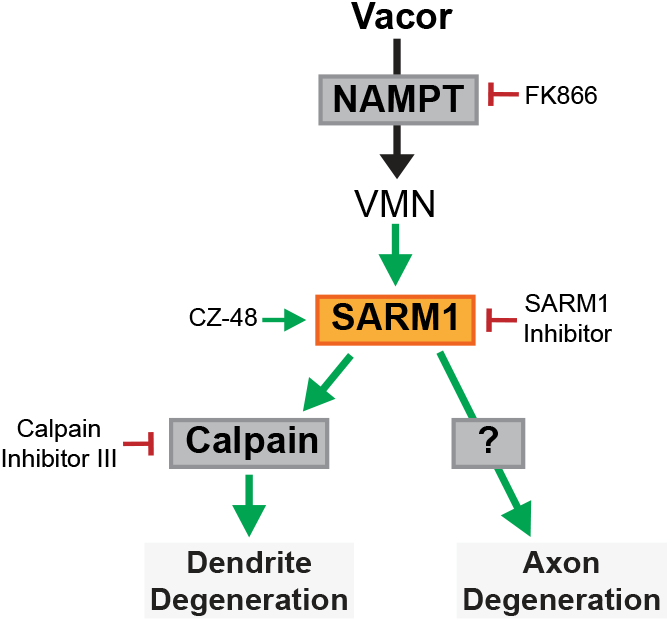
A model showing distinct degenerative pathways in different neuronal compartments in cultured mouse hippocampal neurons.

## Discussion

SARM1 has been studied primarily in axons since its identification as a critical mediator of Wallerian degeneration (1, 28, 29). In this study we sought to understand the localization and function of SARM1 in CNS neurons with more complex cellular morphology. We demonstrated that SARM1 is expressed throughout the neuron and can contribute not only to axon degeneration but also to programmed degeneration of the dendrite and the cell body in response to direct activation. Recently identified SARM1 agonists, CZ-48 and Vacor/VMN (20, 23) allowed us to directly activate SARM1 in the absence of other injury or stress and more specifically dissect SARM1-dependent downstream signaling events in distinct subcellular compartments. Treatment of hippocampal neurons with these compounds resulted in SARM1-dependent degeneration of axons, dendrites, and cell bodies (Figure 2). Surprisingly, SARM1 degeneration in the axons of hippocampal neurons was calpain independent, while degeneration of dendrites required Ca^2+^ uptake and calpain activity. Taken together, these observations demonstrate that activation of SARM1 is sufficient to cause destruction of the entire CNS neuron, though the pathways that mediate degeneration downstream of SARM1 are distinct in axons and dendrites.

Previous studies have implicated calpain proteases in the degeneration of DRG axons following lesion or NGF withdrawal, where calpain inhibition robustly prevents axon degeneration (37, 41, 49). In our primary hippocampal neuron models, we also observed a requirement for calpain activity, but this was specific to dendrites and did not impact axon degeneration (Figure 3A-B). Why these proteases are only critical for dendrite and not axon degeneration in these paradigms is still unclear. Calpains have been shown to be distributed to both axons and dendrites and their activity is required for neurite consolidation in hippocampal neurons (63). Our data show that a canonical axon-specific calpain substrate Nf-L is not processed following SARM1 activation, unlike the well processed substrates found in both axons and dendrites (Figure 6). This observation suggests that calpains are not activated in the axonal compartment downstream of SARM1 as is observed in DRG neurons. It is therefore possible that SARM1’s downstream pathways may depend on neuron subtype and the specific factors that they express. Alternatively, calpain activation in hippocampal axon may be dependent on a secondary upstream input that is not present following pure SARM1 activation. However, the observation that hippocampal axons still degenerate in the absence of calpain activity suggests that distinct pathways can drive this process.

Our finding that calcium influx in dendrites via Cav2 channels can be regulated by SARM1 activity provides new insight into the pathways downstream of this NADase that drive degeneration. The impact of calcium influx has been extensively studied in models of axonal lesion, where diverse models such as mouse SCG/DRGs, zebrafish and flies have revealed two distinct phases of calcium influx in Wallerian degeneration (39, 40, 42, 52–54). The first phase is transient and proximal to the axon injury site, while the second is a more widespread terminal phase of calcium influx that shortly precedes degeneration. There is evidence that the WLDS mutant protein (39) or SARM1 deletion (40, 42) can prevent the second phase of calcium influx, but their impact on the first phase remains to be resolved (39, 40, 52–54). We observed a SARM1-dependent calcium influx in response to Vacor activation that only occurred after two hours and was more pronounced at four hours after SARM1 activation (Figure 5). This indicates that SARM1 activity does not contribute to an early influx of calcium in this experimental setup and supports the earlier findings that SARM1 deletion can specifically impact late-phase calcium influx. In Wallerian degeneration of DRG or SGC neurons, various calcium channels have been implicated in this calcium influx (40, 55, 56), where our data indicate that in primary hippocampal neurons the SARM1-dependent calcium influx is mediated by Cav2 as Cilnidipine, but not Nifedipine, blunted SARM1-dependent calcium influx (Figure 5). Although the specific mechanisms by which SARM1 regulates Cav2 activity remains unclear, it is possibly mediated directly by cADPR generated by SARM1 (20, 57, 58), or indirectly by NAD^+^ loss. cADPR is involved in mobilizing downstream calcium fluxes as this metabolite has many roles in TIR/cADPR/calcium signaling defined in other pathways (59–61).

In this study we defined a degenerative role for SARM1 in hippocampal neurons that is localization dependent, with calpains playing a critical role only in dendrites (Figure 7). A key question remains regarding what pathway(s) execute axon degeneration downstream of SARM1 in hippocampal neurons. Our observation that SARM1-dependent axon degeneration in hippocampal neurons is independent of both calpain and caspase proteases indicates that it is driven by distinct pathways that have yet to be defined. In addition, this result suggests that the contribution of calpains to degeneration of specific cellular processes is not uniformly conserved across neuronal subtypes, and the previously observed role in DRG/SCGs may be specific to these sensory neurons given their unique morphology characterized by a single bifurcating axon. It will also be important to understand the contribution of SARM1 to degeneration of CNS neurons in vivo following the complex cellular insults that occur in neurodegenerative disease or following acute injury. Further studies will be needed to fully understand the impact of SARM1 activity and downstream pathway interactions in axons, dendrites and cell bodies of CNS neurons following diverse insults.

## Experimental procedures

Table 1 lists all antibodies and Table 2 list key reagents used.

### Primary hippocampal neurons

Hippocampi were dissected from E16-E18 embryos of C57BL/6, CD-1 (Charles River Laboratories), or Sarm1-deficient mice (B6. 129X1-*Sarm1^tm1Aidi^*/J, 018069, The Jackson Laboratory) in ice-cold Hibernate E solution. Dissected hippocampi were digested with ∼0.1 unit/hippocampus of papain in 1x PBS at 37°C for 10min and then incubated with DNAseI (3000 unit/ml) and HI-FBS (half the volume of papain solution) at RT for 5min. Hippocampi were dissociated by gentle pipetting (∼20 strokes) in the complete NbActiv4 medium (50µl medium per hippocampus). The complete NbActiv4 medium consists of NbActiv4 (500ml) supplemented with Penicillin-Streptomycin (5ml, the final concentration at 100U/ml), Glutamax (5ml), and 5-floro-2-deoxyuridine (final concentration 1µM). Dissociated hippocampi were spun down at 500rcf at 4°C for 3min and the pellet was dissociated in the complete NbActiv4. Live cells were counted using a Cellometer Auto 2000 Cell Viability Counter (Nexcelom), and the cells were plated on PDL-coated 96-well or 384-well plates at the density of 25000 or 12500 cells/well in 100µl or 50µl of the complete NbActiv4. Hippocampal neurons were grown for 8-12 days with half-medium changes every 7 days. Primary hippocampal neurons from *wildtype* or *Sarm1*-deficient mice are hereafter referred to as *wildtype* or *Sarm1KO* neurons.

### Primary dorsal root ganglia (DRG) neuron

Dorsal root ganglia (DRG) were dissected from E12-E14 embryos of CD-1 (Charles River Laboratories) in ice-cold L15 medium. Dissected DRG were trypsinized by resuspending them in Trypsin-EDTA solution with shaking at 300rpm at 37°C for 15min. After trypsinization, DRG were spun down at 1500rcf at room temperature (RT) for 2min, rinsed once with L15 with 10% HI-FBS, dissociated in L15 with 10% HI-FBS, and triturated by slow pipetting (∼20 strokes). Triturated DRG neurons were spun down at 2000rcf at RT for 2min and resuspended in 1ml of the complete DRG medium. The complete DRG medium consists of Neurobasal medium (500ml) supplemented with B27 supplement (10ml), 1M Glucose (20ml), Glutamax (5ml), Penicillin-Streptomycin (10ml, the final concentration: 200U/ml), Nerve growth factor (NGF, the final concentration at 25ng/ml), and 5-floro-2-deoxyuridine (the final concentration at 1µM). The number of viable cells were counted using Cellometer Auto 2000 Cell Viability Counter (Nexcelom). DRG neurons were spun down again at 2000rcf for 2min at RT. Pellet of DRG neurons were resuspended in the appropriate volume of the complete DRG medium so that the concentration of viable cells is 100000/µl. 1.5µl of dissociated DRG neurons were spotted in the middle of wells on 96w plates that are PDL- and laminin-coated. Plates were kept in the tissue-culture incubator (37°C with 5% CO_2_ and 95% humidity) for 10 min, and then the complete DRG medium was slowly added to each well (50µl/well).

### Immunocytochemistry

Primary hippocampal neurons were treated with CZ-48, Vacor, and/or other pharmacological tools in the complete NbActiv4 medium at 37°C and 5% CO_2_ for 8-16h. After treatment, neurons were fixed with 4% paraformaldehyde (PFA) with 4% sucrose (Sigma-Aldrich, S0389) in 1x PBS at RT for 15min. Fixed neurons were washed with 1x PBS once and then blocked in the blocking buffer at RT for at least 1h. The blocking buffer consists of 5% bovine serum albumin (BSA), 0.5% Triton X100 in 1x PBS. After blocking, neurons were incubated with primary antibodies diluted in the blocking buffer at 4°C for at least 16h. Neurons were then washed 3 times with 1x PBS, incubated with secondary antibodies diluted in the blocking buffer at RT for 1h, and then washed again 3 times with 1x PBS.

### Super resolution confocal imaging and quantification of SARM1 localization

Neurons were plated on PDL-coated 96-well plate (Greiner Bio-one, 655946) for super resolution confocal imaging. Immunocytochemistry was done essentially in the same way as described above, except for fixation. For super resolution confocal imaging, cells were fixed with 4% PFA with 4% sucrose in 1x PBS at RT for 10min and then 100% MeOH at 4°C for an additional 5 min for mitochondrial staining. Immunostained cells were imaged with a laser scanning confocal microscope (Leica SP8; Leica Microsystems, Inc), acquired with a 63x/1.4 NA oil objective in LIGHTNING super-resolution mode with a pixel size of 40 nm. Images were then processed using an adaptive processing algorithm. The representative images were generated by three-dimensional reconstruction using Imaris (Bitplane). To obtain accurate mitochondrial 3D structures, we used surface rendering and fluorescence thresholding. Quantification of SARM1 localization within mitochondria was performed using ImageJ2 (National Institutes of Health), by profiling the fluorescence intensity of each channel by Plot_Multicolor4.3 (64).

### Liquid handling (plating, feeding, fixing, and immunostaining neurons on 384-well plates)

Neurons were resuspended in the complete NbActive4 media at a density of 250000 cells/mL. A Dynamic Devices Lynx liquid handler in a sterile cabinet was used to plate cells (12500 cells /well in 50µl media) into PDL-coated 384-well clear-bottom optical plates, with gentle resuspension of cells in the source reservoir prior to transfer to promote even plating density. Cells feeding was performed in the Lynx by removing 50% of the media and gently adding the complete media slowly to avoid cell disruption every 7 days. Immunostaining was performed as described in the “Immunocytochemistry” section, using a BioTek EL406 plate washer and Agilent Bravo liquid handler, integrated by HighRes Biosolutions, to gently transfer 1x PBS or reagents to neurons at each step.

### Quantification of axon/dendrite degeneration and nuclear condensation

Immunostained neurons were imaged using the Opera Phenix High Content Screening System (PerkinElmer) and images were quantified using the Harmony software (PerkinElmer). To acquire fluorescent images, we used built-in acquisition protocols for DAPI (excitation/emission at 375nm/ 435-480nm), Alexa488 (488nm/ 500-550nm), Alexa568 (561nm/ 570-630nm), and Alexa647 (640nm/ 650-760nm), and water-immersion 40x objective (NA=1.1) using the confocal mode acquiring two z-slice images spaced 1µm apart. Areas of axons or dendrites were quantified by applying the built-in spot analysis on images of respective channels. Total areas of axons or dendrites were quantified per well and per plate, and normalized to the respective control values (e.g. mean of untreated wildtype wells) per plate. All the nuclei were detected by applying the built-in nucleus detection protocol on DAPI images and further classified into condensed or normal nuclei by applying a manual linear classifier that was trained using at least 50 nuclei per group.

### Quantification of PAD6 signal

Cultured neurons were incubated with PC6 (50µM) and CellMask deep red (Thermo Fisher Scientific C10046, 1:5000) for 2h prior to imaging, in the complete NbActiv4 in the humidified incubator at 37°C with 5% CO2. 1h prior to imaging, SARM1 agonist and inhibitors were added by D300e digital dispenser (Tecan) and further incubated in the incubator. PAD6 signal was imaged using the Opera Phenix Plus High Content Screening System (PerkinElmer) and images were quantified using the Harmony software (PerkinElmer). To acquire PAD6 and CellMask signal, we used built-in acquisition protocols for mTurquoise2-extended (excitation/emission at 405nm/435-550nm) and Alexa647 (640nm/ 650-760nm), respectively, and water immersion 63x (NA=1.15) under the confocal mode acquiring two z-slice images spaced 0.5µm apart, at 37°C and with 5% CO_2_. PAD6-positive area was quantified first by applying an intensity threshold for mTurquoise2-extended channel to select pixels with PAD6 signal above the background, and then by measuring the total numbers of pixels per well. The PAD6-positive area was normalized to the respective control values (mean of untreated wildtype wells) per plate.

### Calcium imaging using Fluo4

*Wildtype* or *Sarm1KO* neurons were incubated with Fluo4 (1µM) in for 30min prior to the compound addition, in the complete NbActiv4 in the humidified incubator at 37°C with 5% CO2. SARM1 agonist and inhibitors were added by D300e digital dispenser (Tecan) and imaging was started immediately. Fluo4 signal was imaged using the Opera Phenix Plus High Content Screening System (Perkin Elmer) and images were quantified using the Harmony software (PerkinElmer). To acquire Fluo4 signal, we used built-in acquisition protocol for Fluo4 (excitation/emission at 488nm/500-550nm) and water immersion 20x (NA=1.0) under the confocal mode at 37°C with 5% CO2. Fluo4 signal intensity was quantified first by applying the built-in spot analysis to define Fluo4-positive puncta. Secondly, corrected spot intensity of Fluo4 signal was integrated per each spot and then summarized per well at each time point. The summarized Fluo4 intensity was normalized to the mean intensity of the vehicle-treated well at the first time point (t=0min after addition of toxins and inhibitors) in each genotype per biological replicate

### Western blot

Equal numbers of cultured neurons underwent treatment, were lysed in equal volumes of lysis buffer (1x NuPAGE LDS sample buffer and 1x NuPAGE Sample Reducing Agent), were denatured at 95°C for 10min, and were loaded per gel lane. Protein samples were electrophoresed on NuPAGE Novex 4-12% Bis-Tris Midi protein gels in 1x NuPAGE MOPS SDS running buffer at 200V for 1h at RT. Gels were transferred to nitrocellulose membranes with Tran-Blot Turbo Transfer System (Bio-Rad). Membranes were blocked with Odyssey blocking buffer (Li-Cor 927-40000) at RT for at least 1h, incubated with primary antibodies overnight at 4°C followed by 4x washes with TBST at RT, and then incubated with secondary antibodies at RT for 1h, followed by 4x washes with TBST at RT. Li-Cor odyssey system was used for western blot detection and quantification. To quantify the calpain activity (Figure 6), the signal intensities of full-length and cleaved substrates were separately quantified.

### NAD(H) quantification

Equal numbers of *wildtype* or *Sarm1KO* neurons plated on 96-well plates were treated with various concentrations of CZ-48, or Vacor for 16 hours. Cellular NAD(H) levels were measured using NAD/NADH-Glo assay (Promega). Briefly, neuron plates were cooled down to RT for 30min and the medium volumes were normalized to 50µl per well. The NAD/NADH-Glo detection reagent was prepared while neuron plates were at RT and applied to neuron plates (50µl/well). Plates were incubated at RT for 1h with gentle horizontal shake at 300rpm. Luminescence of each well was measured using EnVision Multimode Plate Reader (PerkinElmer).

### Synthesis of CZ-48, PC6, and SARM1 inhibitor

CZ-48 (65), PC6 (35), and SARM1 inhibitor 3-iodo-1H-pyrrolo[2,3-c]pyridine (36) were synthesized by following the methods described previously.

### Mouse Handling

All mouse procedures adhered to regulations and protocols approved by Denali Therapeutics Institutional Animal Care and Use Committee. Mice were housed under a 12-hour light/dark cycle and had access to water and standard rodent diet *ad libitum*.

### Experimental Design, Data Collection, and Statistical Analysis

Each experiment consists of all the conditions that were directly compared, including genotype, variable doses of toxin treatment, and the presence of pharmacological treatment. Each condition consists of 1-3 technical replicates, such as wells in western blot and immunocytochemistry. Bulk of data were acquired and analyzed by automated systems in an unbiased manner. Super-resolution microscopy (Figures 1A, Figure Supplement 6-1) and western blot (Figure 6) experiments were performed and analyzed in a non-blinded manner. Acquired data were averaged and/or normalized to the control condition (e.g. DMSO-treated wildtype neurons) in each experiment. We then repeated experiments using biologically distinct samples, which indicate neurons prepared from different mice on a different day (typically one to several weeks apart). Averaged and/or normalized values from each biologically independent replicate was treated as independent unit (n), plotted, and reported in each Figure legend.

Statistical tests were performed by ANOVA using stats::aov function in R (Version 4.3.0) and independent variables were reported in Figure legend. An interaction effect between two independent variables was also assessed when running two-way ANOVA. When ANOVA indicated a significant (P < 0.05) effect of an independent variable of interest (e.g. genotype and pharmacological treatment), either Tukey’s (if numbers of groups were 3 or less) or Dunnett’s (4 or more) post hoc test was further performed to compare specific points (e.g. the difference between two genotypes at a specific toxin dose) using pairs or contrast function from the emmeans R package (66). P values that are less than 0.05 are reported in Figure panels using asterisks that represent the ranges of P values – ns, *, **, and *** indicate P ≥ 0.05, P < 0.05, P < 0.01, and P < 0.001 respectively.

### Data availability

The data described in the manuscript are contained within the manuscript as graphs. Raw data can be shared upon request – please contact the corresponding authors.

## Acknowledgements

We thank Thomas Sandmann for statistical analysis support and manuscript feedback. We also thank members of Denali Therapeutics for helpful discussions and comments on the manuscript. This study was funded by Denali Therapeutics.

---

**Figure Supplement 1-1.**
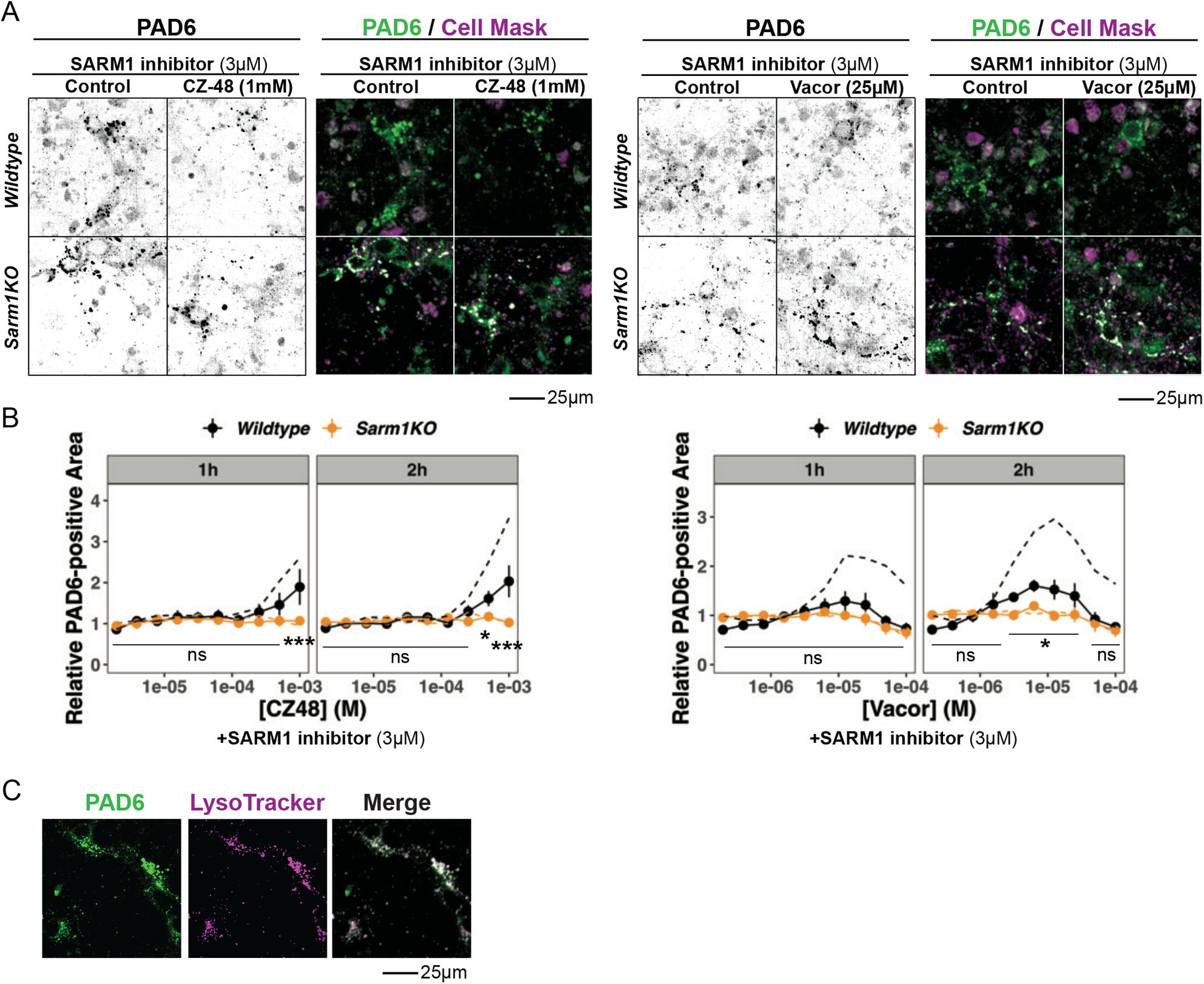
SARM1 inhibitor treatment prevent PAD6 increases following SARM1 activation. (**A**) Representative images following 1 hour treatment and (**B**) quantification of PAD6 in *wildtype* and *Sarm1KO* neurons treated with CZ-48 or Vacor for 1 or 2 hours. Neurons were also co-treated with SARM1 inhibitor (3 µM). For reference, the data from Figure 1D and F (the same experiment without SARM1 inhibitor) are shown in dashed lines. (**C**) Representative images of *wildtype* neurons that were loaded with PC6 and LysoTracker Deep Red for 20 min and imaged for the fluorescent channels for PAD6 and LysoTracker. In (**B**), statistical significance of differences was determined by two-way ANOVA using genotype and toxin doses as two independent variables. When the effect of genotype was significant (P < 0.05), Tukey’s post hoc test was performed and P values that are less than 0.05 for comparisons between the two genotypes at the specific toxin doses are shown. Numbers of biologically independent experiments (n) are 3. Data represent the mean ± SEM, * P < 0.05, ** P < 0.01, *** P < 0.001, ns P ≥ 0.05.

**Figure Supplement 2-1.**
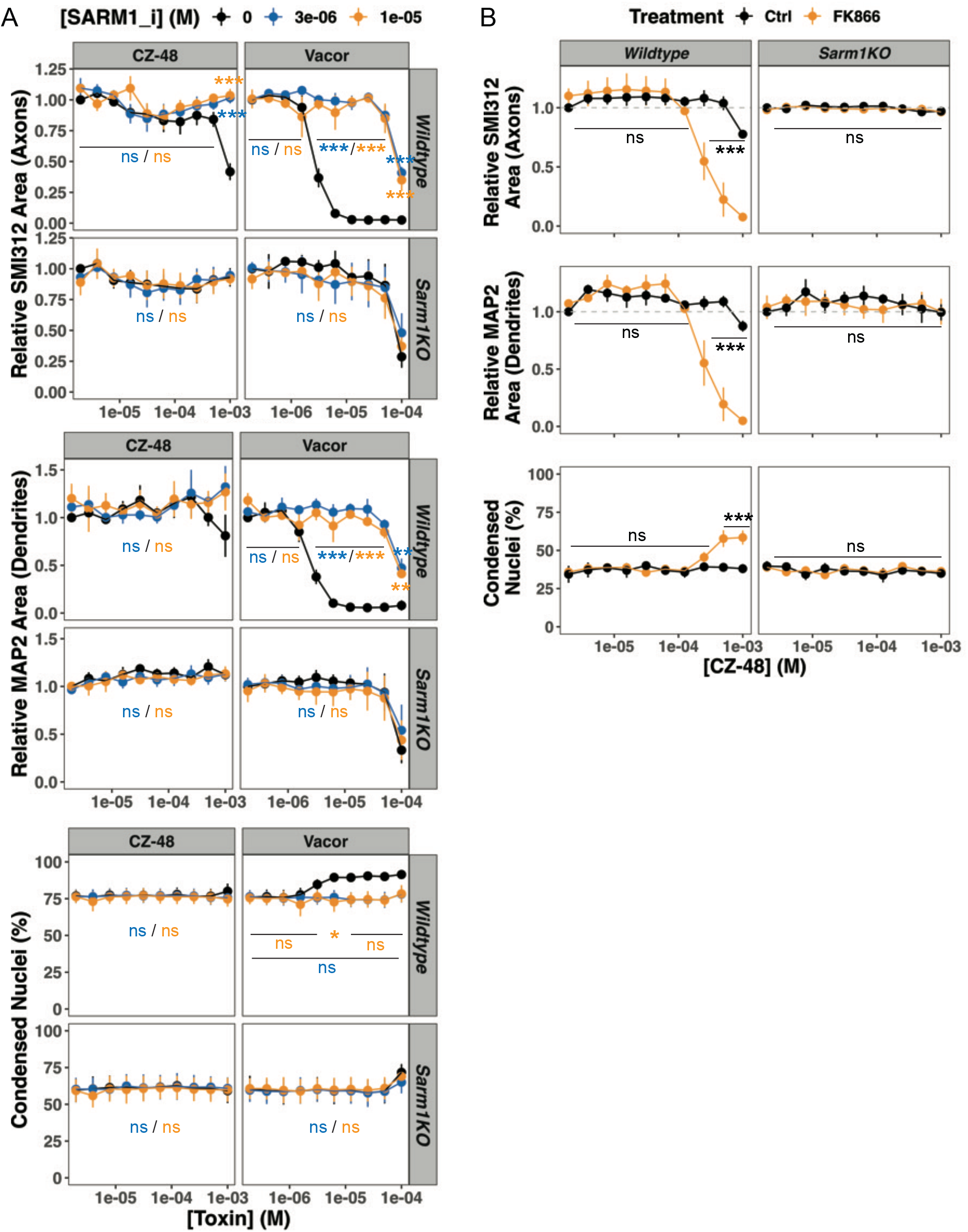
SARM1 inhibitor protects neuronal degeneration induced by SARM1 agonists and FK866 enhances neuronal degeneration induced by CZ-48. Relative levels of immunostained axons and dendrites, and proportion of condensed nuclei in *wildtype* and *Sarm1KO* neurons that were treated with increasing concentrations of (**A**) CZ-48 or Vacor in the presence of the SARM1 inhibitor (SARM1_i: 0, 3, or 10µM) for 16 hours, or (**B**) CZ-48 in the presence of the NAMPT inhibitor FK866 (100nM) for 8 hours. Statistical significance was determined by two-way ANOVA using (**A**) SARM1_i treatment and toxin doses or (**B**) FK866 treatment and toxin doses as two independent variables. When the effect of the former independent variable was significant (P < 0.05), Tukey’s post hoc test was performed and P values that are less than 0.05 for the effect of (**A**) SARM1_i doses relative to the control (black lines) or (**B**) FK866 treatment at the specific toxin doses are shown in figure panels. In (**A**), the ranges of P values are shown in a color-matched manner. Numbers of biologically independent experiments (n) are (**A**) 3-4 and (**B**) 3. Data represent the mean ± SEM, * P < 0.05, ** P < 0.01, *** P < 0.001, ns P ≥ 0.05.

**Figure Supplement 3-1.**
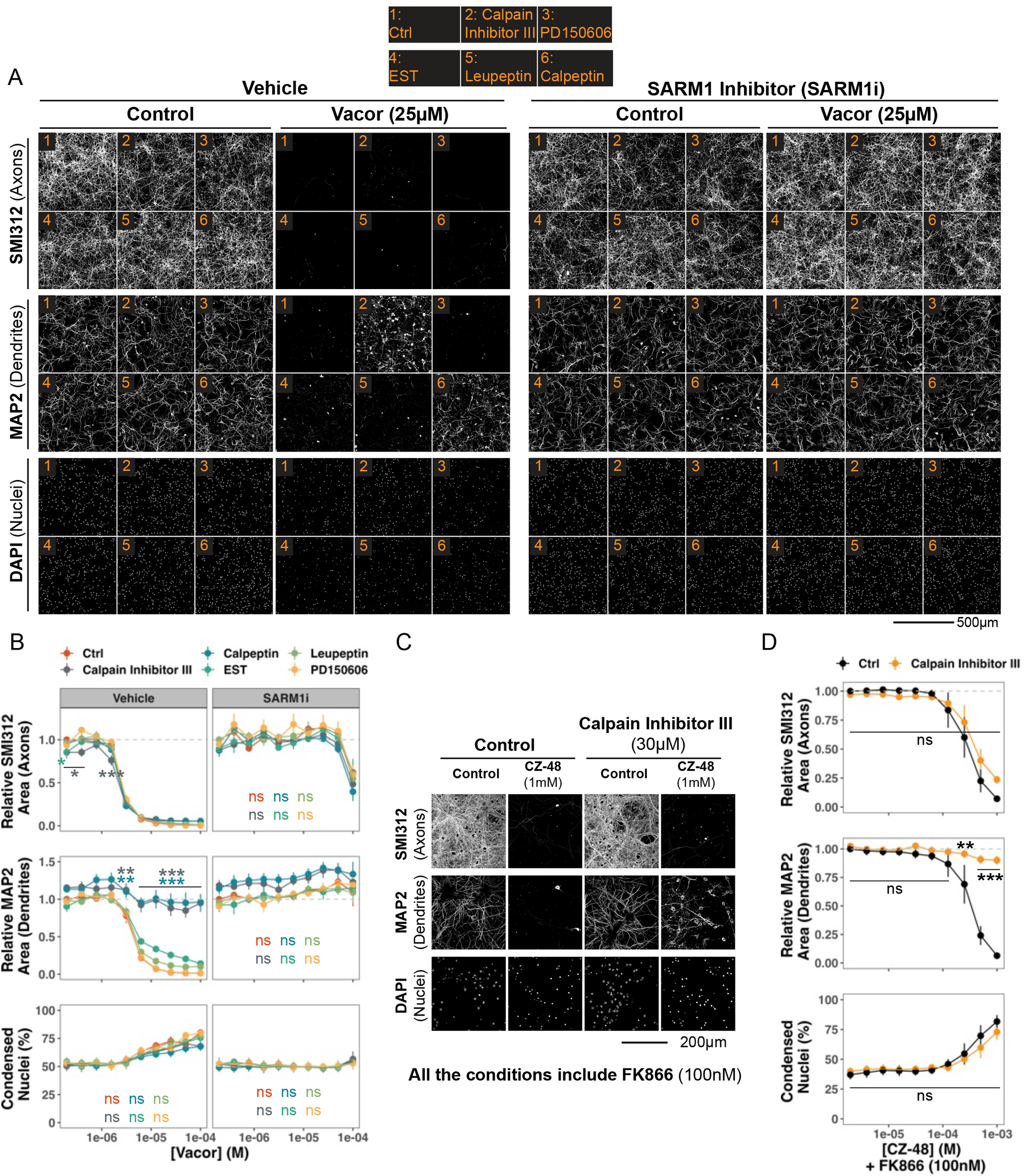
Calpains are required for SARM1-induced dendrite degeneration. (**A**) Representative images of *wildtype* neurons treated with increasing concentrations of Vacor for 16 hours along with the indicated calpain inhibitors listed in the legend on top of images (all 30µM). (**B**) Quantification of relative levels of immunostained axons and dendrites, and proportion of condensed nuclei in (**A**). (**C**) Representative images of *wildtype* neurons co-treated with increasing concentrations of CZ-48 and 100nM FK866 for 8 hours with or without Calpain Inhibitor III (30µM). (**D**) Quantification of relative levels of immunostained axons and dendrites, and proportion of condensed nuclei in (**C**). Statistical significance of effects by calpain inhibitors was determined by two-way ANOVA using inhibitor treatment and toxin ((**B**) Vacor or (**D**) CZ-48) doses as two independent variables. When the effect of the inhibitor treatment was significant (P < 0.05), either (**B**) Dunnett’s or (**D**) Tukey’s post hoc test was performed and P values that are less than 0.05 for the effect of inhibitor treatment relative to the control condition at the specific toxin doses are shown in figure panels. In (**B**), the ranges of P values are shown in a color-matched manner. Numbers of biologically independent experiments (n) are (**B**) 4 and (**D**) 5. Data represent the mean ± SEM, * P < 0.05, ** P < 0.01, *** P < 0.001, ns P ≥ 0.05.

**Figure Supplement 3-2.**
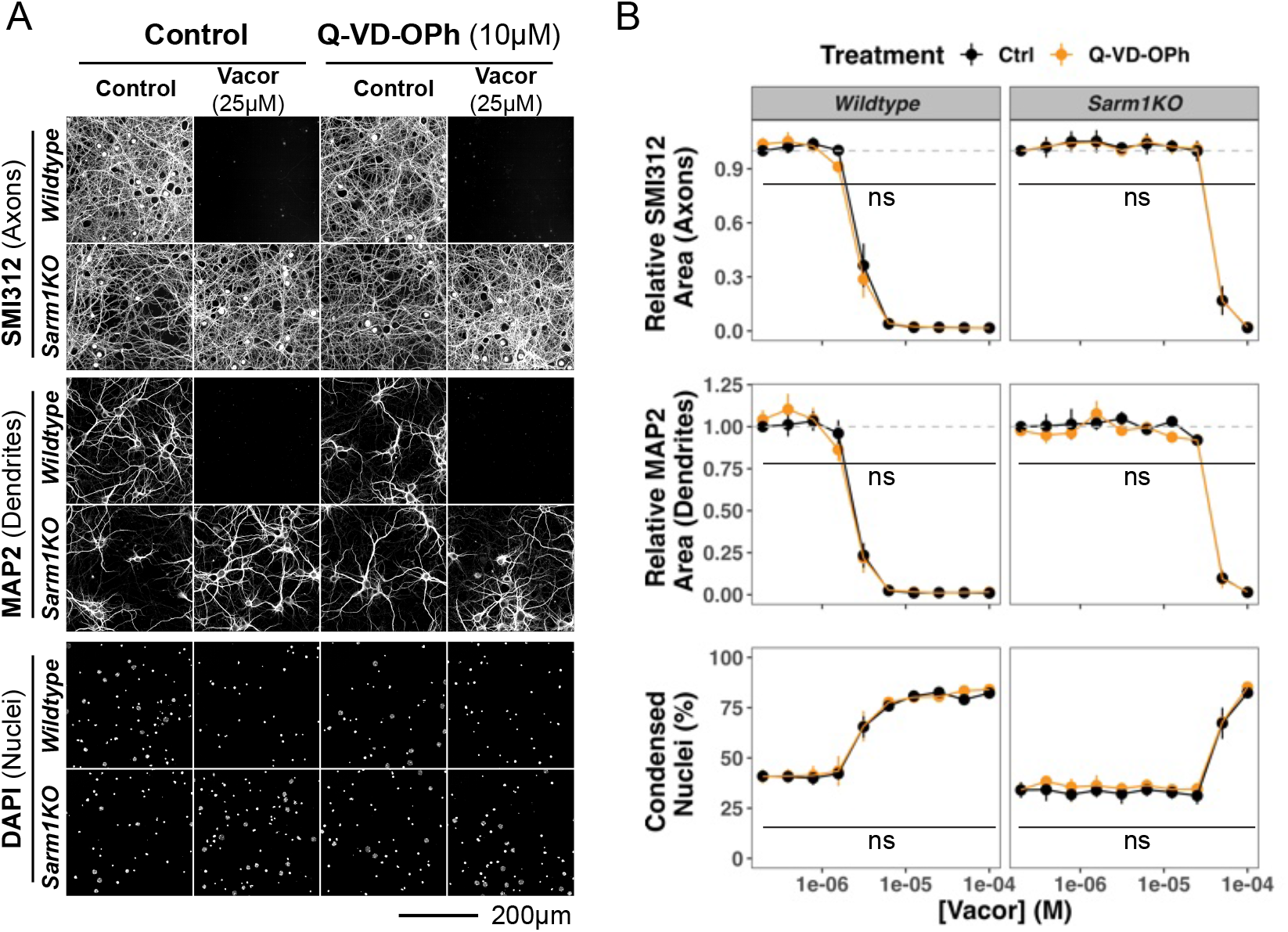
Caspases are not required for SARM1-induced degeneration. (**A**) Representative images and (**B**) relative levels of immunostained axons and dendrites, and proportion of condensed nuclei in *wildtype* and *Sarm1KO* neurons that were treated with increasing concentrations of Vacor for 16 hours with or without the pan-caspase inhibitor Q-VD-OPh (10µM). Statistical significance of effects by Q-VD-OPh treatment was determined by two-way ANOVA using Q-VD-OPh treatment and Vacor doses as two independent variables. The effect of Q-VD-OPh treatment was not significant (P ≥ 0.05) in any readouts. Numbers of biologically independent experiments (n) are 3. Data represent the mean ± SEM, ns P ≥ 0.05.

**Figure Supplement 5-1.**
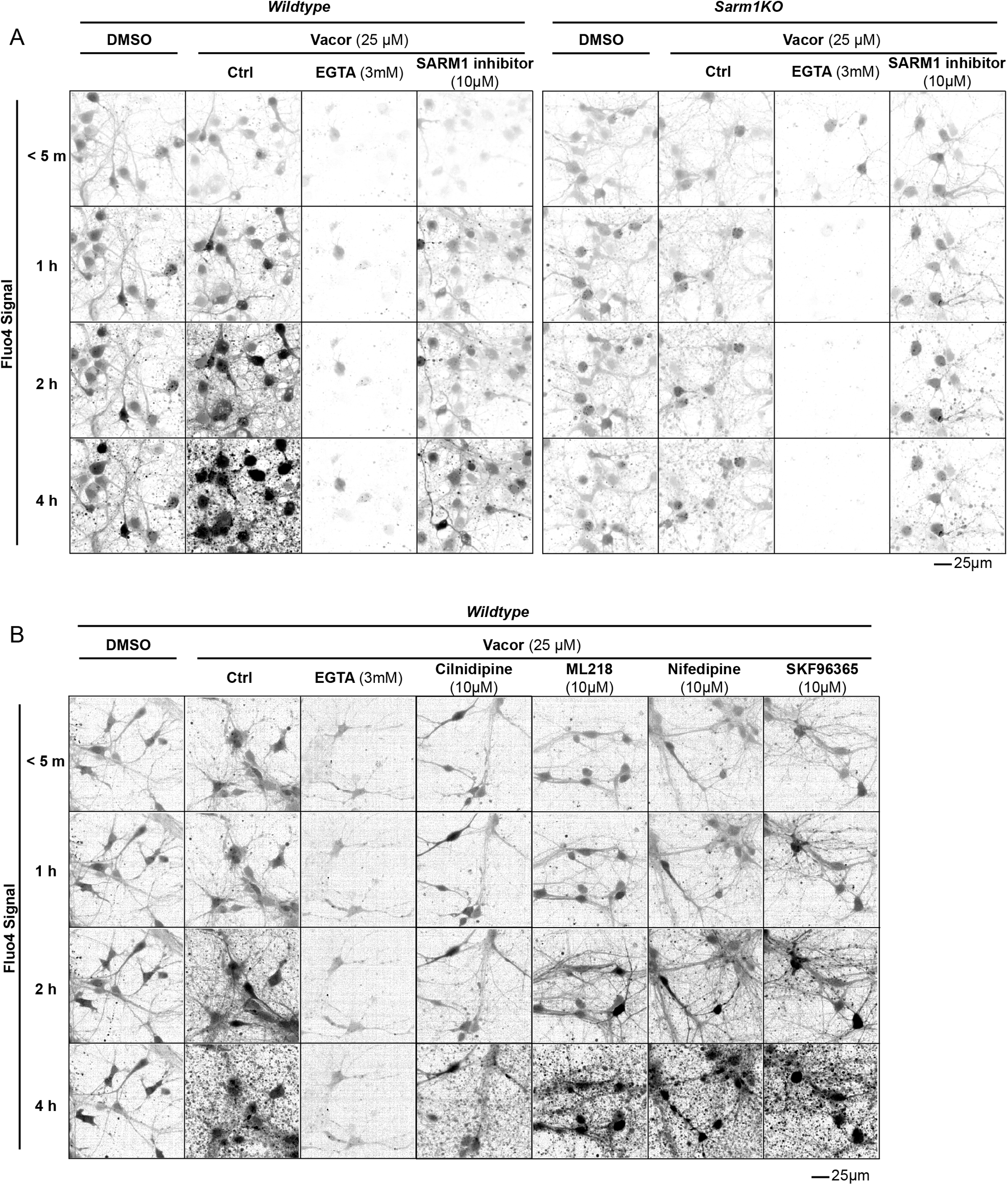
SARM1 activation increases intracellular Ca^2+^. Representative Fluo4 images of *wildtype* or *Sarm1KO* neurons that were treated with Vacor (25µM) or DMSO for < 5min, 1, 2, and 4 hours. (**A**) Neurons were also co-treated with EGTA (3mM), SARM1 inhibitor (10µM) or (**B**) indicated Cav channel blockers (all at 10µM).

**Figure Supplement 6-1.**
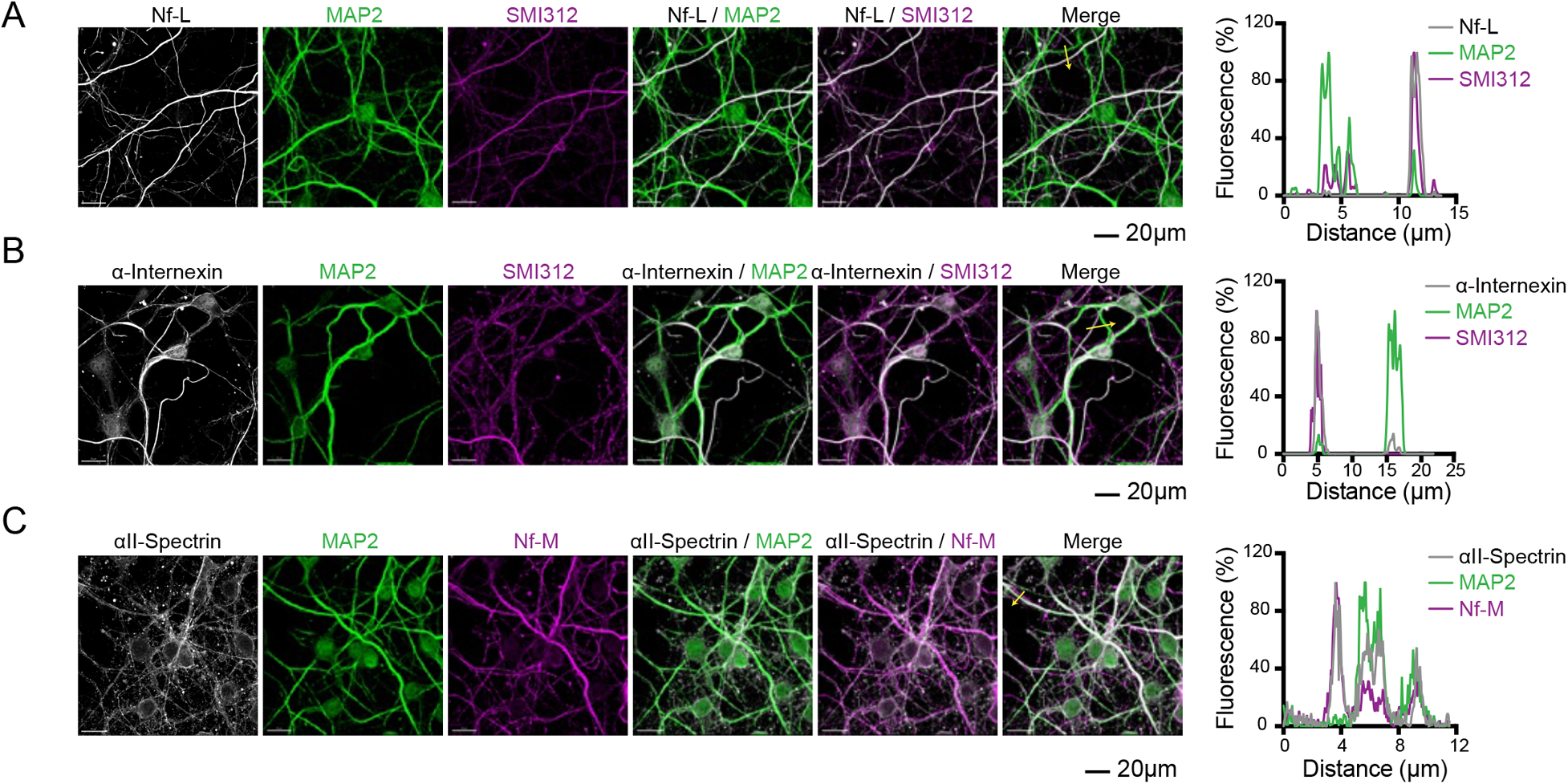
Localization of Neurofilament light, α-Internexin, and αII-Spectrin in axons and dendrites in primary mouse hippocampal neurons. Representative immunofluorescent images and line graphs of primary mouse hippocampal neurons from *wildtype* mice. Neurons were immunostained for (**A**) Neurofilament light (Nf-L), (**B**) α-Internexin or (**C**) αII-Spectrin along with axonal (SMI312 or Neurofilament middle, Nf-M) and dendritic (MAP2) marker proteins. Line graphs on the right indicate the relative intensity of the immunostaining along the yellow arrows indicated in the merged images.

## References

1. Coleman, M. P., and Höke, A. (2020) Programmed axon degeneration: from mouse to mechanism to medicine. Nat. Rev. Neurosci. 21, 183–196

2. Mcguinness, H. Y., Gu, W., Shi, Y., Kobe, B., and Ve, T. (2023) SARM1-Dependent Axon DegenerationL: Nucleotide Signaling, Neurodegenerative Disorders, Toxicity, and Therapeutic Opportunities. 10.1177/10738584231162508

3. Lunn, E. R., Perry, V. H., Brown, M. C., Rosen, H., and Gordon, S. (1989) Absence of Wallerian degeneration does not hinder regeneration in peripheral nerve. Eur. J. Neurosci. 1, 27–33

4. Mack, T. G., Reiner, M., Beirowski, B., Mi, W., Emanuelli, M., Wagner, D., Thomson, D., Gillingwater, T., Court, F., Conforti, L., Fernando, F. S., Tarlton, A., Andressen, C., Addicks, K., Magni, G., Ribchester, R. R., Perry, V. H., and Coleman, M. P. (2001) Wallerian degeneration of injured axons and synapses is delayed by a Ube4b/Nmnat chimeric gene. Nat. Neurosci. 4, 1199–1206

5. Araki, T., Sasaki, Y., and Milbrandt, J. (2004) Increased Nuclear NAD Biosynthesis and SIRT1 Activation Prevent Axonal Degeneration. Science. 305, 1010–1013

6. Conforti, L., Wilbrey, A., Morreale, G., Janeckova, L., Beirowski, B., Adalbert, R., Mazzola, F., Stefano, M. Di, Hartley, R., Babetto, E., Smith, T., Gilley, J., Billington, R. A., Genazzani, A. A., Ribchester, R. R., Magni, G., and Coleman, M. (2009) Wld S protein requires Nmnat activity and a short N-terminal sequence to protect axons in mice. J. Cell Biol. 184, 491–500

7. Babetto, E., Beirowski, B., Janeckova, L., Brown, R., Gilley, J., Thomson, D., Ribchester, R. R., and Coleman, M. P. (2010) Targeting NMNAT1 to axons and synapses transforms its neuroprotective potency in vivo. J. Neurosci. 30, 13291–13304

8. Sasaki, Y., Vohra, B. P. S., Baloh, R. H., and Milbrandt, J. (2009) Transgenic mice expressing the Nmnat1 protein manifest robust delay in axonal degeneration in vivo. J. Neurosci. 29, 6526–6534

9. Gilley, J., and Coleman, M. P. (2010) Endogenous Nmnat2 Is an Essential Survival Factor for Maintenance of Healthy Axons. PLoS Biol. 8, e1000300

10. Osterloh, J. M., Yang, J., Rooney, T. M., Fox, A. N., Adalbert, R., Powell, E. H., Sheehan, A. E., Avery, M. a, Hackett, R., Logan, M. a, MacDonald, J. M., Ziegenfuss, J. S., Milde, S., Hou, Y.-J., Nathan, C., Ding, A., Brown, R. H., Conforti, L., Coleman, M., Tessier-Lavigne, M., Züchner, S., and Freeman, M. R. (2012) dSarm/Sarm1 is required for activation of an injury-induced axon death pathway. Science. 337, 481–4

11. Essuman, K., Summers, D. W., Sasaki, Y., Mao, X., Diantonio, A., and Milbrandt, J. (2017) The SARM1 Toll / Interleukin-1 receptor domain possesses intrinsic NAD^+^ cleavage activity that promotes pathological axonal degeneration. Neuron. 93, 1334–1343.e5

12. Loring, H. S., Icso, J. D., Nemmara, V., and Thompson, P. R. (2020) Initial kinetic characterization of sterile alpha and toll/interleukin receptor motif-containing protein 1. Biochemistry. 59, 933–942

13. Horsefield, S., Burdett, H., Zhang, X., Manik, M. K., Shi, Y., Chen, J., Qi, T., Gilley, J., Lai, J. S., Rank, M. X., Casey, L. W., Gu, W., Ericsson, D. J., Foley, G., Hughes, R. O., Bosanac, T., Von Itzstein, M., Rathjen, J. P., Nanson, J. D., Boden, M., Dry, I. B., Williams, S. J., Staskawicz, B. J., Coleman, M. P., Ve, T., Dodds, P. N., and Kobe, B. (2019) NAD+ cleavage activity by animal and plant TIR domains in cell death pathways. Science. 365, 793–799

14. Sporny, M., Guez-Haddad, J., Khazma, T., Yaron, A., Dessau, M., Shkolnisky, Y., Mim, C., Isupov, M. N., Zalk, R., Hons, M., and Opatowsky, Y. (2020) Structural basis for SARM1 inhibition and activation under energetic stress. eLife. 9, 1–25

15. Di Stefano, M., Nascimento-Ferreira, I., Orsomando, G., Mori, V., Gilley, J., Brown, R., Janeckova, L., Vargas, M. E., Worrell, L. A., Loreto, A., Tickle, J., Patrick, J., Webster, J. R. M., Marangoni, M., Carpi, F. M., Pucciarelli, S., Rossi, F., Meng, W., Sagasti, A., Ribchester, R. R., Magni, G., Coleman, M. P., and Conforti, L. (2015) A rise in NAD precursor nicotinamide mononucleotide (NMN) after injury promotes axon degeneration. Cell Death Differ. 22, 731–742

16. Alexandris, A. S., Ryu, J., Rajbhandari, L., Harlan, R., McKenney, J., Wang, Y., Aja, S., Graham, D., Venkatesan, A., and Koliatsos, V. E. (2022) Protective effects of NAMPT or MAPK inhibitors and NaR on Wallerian degeneration of mammalian axons. Neurobiol. Dis. 171, 105808

17. Sasaki, Y., Nakagawa, T., Mao, X., DiAntonio, A., and Milbrandt, J. (2016) NMNAT1 inhibits axon degeneration via blockade of SARM1-mediated NAD ^+^ depletion. eLife. 5, 1–15

18. Figley, M. D., Gu, W., Nanson, J. D., Shi, Y., Sasaki, Y., Cunnea, K., Malde, A. K., Jia, X., Luo, Z., Saikot, F. K., Mosaiab, T., Masic, V., Holt, S., Hartley-Tassell, L., McGuinness, H. Y., Manik, M. K., Bosanac, T., Landsberg, M. J., Kerry, P. S., Mobli, M., Hughes, R. O., Milbrandt, J., Kobe, B., DiAntonio, A., and Ve, T. (2021) SARM1 is a metabolic sensor activated by an increased NMN/NAD+ ratio to trigger axon degeneration. Neuron. 109, 1118–1136.e11

19. Loreto, A., Hill, C. S., Hewitt, V. L., Orsomando, G., Angeletti, C., Gilley, J., Lucci, C., Sanchez-Martinez, A., Whitworth, A. J., Conforti, L., Dajas-Bailador, F., and Coleman, M. P. (2020) Mitochondrial impairment activates the Wallerian pathway through depletion of NMNAT2 leading to SARM1-dependent axon degeneration. Neurobiol. Dis. 134, 104678

20. Zhao, Z. Y., Xie, X. J., Li, W. H., Liu, J., Chen, Z., Zhang, B., Li, T., Li, S. L., Lu, J. G., Zhang, L., Zhang, L. he, Xu, Z., Lee, H. C., and Zhao, Y. J. (2019) A cell-permeant mimetic of NMN activates SARM1 to produce cyclic ADP-ribose and induce non-apoptotic cell death. iScience. 15, 452–466

21. Bratkowski, M., Xie, T., Thayer, D. A., Lad, S., Mathur, P., Yang, Y. S., Danko, G., Burdett, T. C., Danao, J., Cantor, A., Kozak, J. A., Brown, S. P., Bai, X., and Sambashivan, S. (2020) Structural and mechanistic regulation of the pro-degenerative NAD hydrolase SARM1. Cell Rep. 10.1016/j.celrep.2020.107999

22. Jiang, Y., Liu, T., Lee, C. H., Chang, Q., Yang, J., and Zhang, Z. (2020) The NAD^+^-mediated self-inhibition mechanism of pro-neurodegenerative SARM1. Nature. 588, 658–663

23. Loreto, A., Angeletti, C., Gu, W., Osborne, A., Nieuwenhuis, B., Gilley, J., Merlini, E., Arthur-Farraj, P., Amici, A., Luo, Z., Hartley-Tassell, L., Ve, T., Desrochers, L. M., Wang, Q., Kobe, B., Orsomando, G., and Coleman, M. P. (2021) Neurotoxin-mediated potent activation of the axon degeneration regulator SARM1. eLife. 10, 1–24

24. Loreto, A., Antoniou, C., Merlini, E., Gilley, J., and Coleman, M. P. (2023) NMN: The NAD precursor at the intersection between axon degeneration and anti-ageing therapies. Neurosci. Res. 10.1016/j.neures.2023.01.004

25. Angeletti, C., Amici, A., Gilley, J., Loreto, A., Trapanotto, A. G., Antoniou, C., Merlini, E., Coleman, M. P., and Orsomando, G. (2022) SARM1 is a multi-functional NAD(P)ase with prominent base exchange activity, all regulated bymultiple physiologically relevant NAD metabolites. iScience. 25, 103812

26. Nan Hou, Y., Cai, Y., Hua Li, W., Ming He, W., Ying Zhao, Z., Jie Zhu, W., Wang, Q., Liu, J., Cheung Lee, H., Goran, S., Zhang, H., and Juan Zhao, Y. (2022) A Conformation-specific Nanobody Targeting the NMN-activated State of SARM1. Nat. Commun. 13, 7898

27. Icso, J. D., and Thompson, P. R. (2022) The chemical biology of NAD+ regulation in axon degeneration. Curr. Opin. Chem. Biol. 69, 102176

28. Krauss, R., Bosanac, T., Devraj, R., Engber, T., and Hughes, R. O. (2020) Axons matter: the promise of treating neurodegenerative disorders by targeting SARM1-mediated axonal degeneration. Trends Pharmacol. Sci. 41, 281–293

29. Gerdts, J., Summers, D. W., Milbrandt, J., and DiAntonio, A. (2016) Axon self-destruction: new links among SARM1, MAPKs, and NAD+ metabolism. Neuron. 89, 449–460

30. Tao, J., and Rolls, M. M. (2011) Dendrites have a rapid program of injury-induced degeneration that is molecularly distinct from developmental pruning. J. Neurosci. 31, 5398–5405

31. Ji, H., Sapar, M. L., Sarkar, A., Wang, B., and Han, C. (2022) Phagocytosis and self-destruction break down dendrites of Drosophila sensory neurons at distinct steps of Wallerian degeneration. Proc. Natl. Acad. Sci. U. S. A. 119, 1–12

32. Buonvicino, D., Mazzola, F., Zamporlini, F., Resta, F., Ranieri, G., Camaioni, E., Muzzi, M., Zecchi, R., Pieraccini, G., Dölle, C., Calamante, M., Bartolucci, G., Ziegler, M., Stecca, B., Raffaelli, N., and Chiarugi, A. (2018) Identification of the nicotinamide salvage pathway as a new toxification route for antimetabolites. Cell Chem. Biol. 25, 471–482.e7

33. LeWitt, P. (1980) The neurotoxicity of the rat poison vacor. A clinical study of 12 cases. N. Engl. J. Med. 302, 73–7

34. Gallanosa, A. G., Spyker, D. A., and Curnow, R. T. (1981) Diabetes mellitus associated with autonomic and peripheral neuropathy after vacor rodenticide poisoning: A review. Clin. Toxicol. 18, 441–449

35. Li, W. H., Huang, K., Cai, Y., Wang, Q. W., Zhu, W. J., Hou, Y. N., Wang, S., Cao, S., Zhao, Z. Y., Xie, X. J., Du, Y., Lee, C. S., Lee, H. C., Zhang, H., and Zhao, Y. J. (2021) Permeant fluorescent probes visualize the activation of SARM1 and uncover an anti-neurodegenerative drug candidate. eLife. 10, 1–21

36. 36. Hughes, R. O., Devraj, R., Bosanac, T., Jarjes-pike, R. A., Brearley, A. S., and Bentley, J. (2019) Inhibitors of Sarm1

37. Yang, J., Weimer, R. M., Kallop, D., Olsen, O., Wu, Z., Renier, N., Uryu, K., and Tessier-Lavigne, M. (2013) Regulation of axon degeneration after injury and in development by the endogenous calpain inhibitor calpastatin. Neuron. 80, 1175–1189

38. Summers, D. W., DiAntonio, A., and Milbrandt, J. (2014) Mitochondrial dysfunction induces Sarm1-dependent cell death in sensory neurons. J. Neurosci. 34, 9338–9350

39. Vargas, M. E., Yamagishi, Y., Tessier-Lavigne, M., and Sagasti, A. (2015) Live imaging of calcium dynamics during axon degeneration reveals two functionally distinct phases of calcium influx. J. Neurosci. 35, 15026–15038

40. Loreto, A., Di Stefano, M., Gering, M., and Conforti, L. (2015) Wallerian degeneration Is executed by an NMN-SARM1-dependent late Ca2+ Influx but only modestly influenced by mitochondria. Cell Rep. 13, 2539–2552

41. Cetinkaya-Fisgin, A., Luan, X., Reed, N., Jeong, Y. E., Oh, B. C., and Hoke, A. (2020) Cisplatin induced neurotoxicity is mediated by Sarm1 and calpain activation. Sci. Rep. 10, 1–12

42. Ko, K. W., Devault, L., Sasaki, Y., Milbrandt, J., and Diantonio, A. (2021) Live imaging reveals the cellular events downstream of sarm1 activation. eLife. 10, 1–21

43. Ono, Y., Saido, T. C., and Sorimachi, H. (2016) Calpain research for drug discovery: Challenges and potential. Nat. Rev. Drug Discov. 15, 854–876

44. Ennes-Vidal, V., Menna-Barreto, R. F. S., Branquinha, M. H., Dos Santos, A. L. S., and D’Avila-Levy, C. M. (2017) Why calpain inhibitors are interesting leading compounds to search for new therapeutic options to treat leishmaniasis? Parasitology. 144, 117–123

45. Tsujinaka, T., Kajiwara, Y., Kambayashi, J., Sakon, M., Higuchi, N., Tanaka, T., and Mori, T. (1988) Synthesis of a new cell penetrating calpain inhibitor (calpeptin). Biochem. Biophys. Res. Commun. 153, 1201–1208

46. Simon, D. J., Weimer, R. M., Mclaughlin, T., Kallop, D., Stanger, K., Yang, J., O’Leary, D. D. M., Hannoush, R. N., and Tessier-Lavigne, M. (2012) A caspase cascade regulating developmental axon degeneration. J. Neurosci. 32, 17540–17553

47. Gerdts, J., Summers, D. W., Sasaki, Y., DiAntonio, A., and Milbrandt, J. (2013) Sarm1-mediated axon degeneration requires both SAM and TIR interactions. J. Neurosci. 33, 13569–80

48. Ozawa, Y., Hayashi, K., and Kobori, H. (2006) New Generation Calcium Channel Blockers in Hypertensive Treatment. Curr. Hypertens. Rev. 2, 103–111

49. Ma, M., Ferguson, T. A., Schoch, K. M., Li, J., Qian, Y., Shofer, F. S., Saatman, K. E., and Neumar, R. W. (2013) Calpains mediate axonal cytoskeleton disintegration during Wallerian degeneration. Neurobiol Dis. 56, 34–46

50. Cryns, V. L., Bergeron, L., Zhu, H., Li, H., and Yuan, J. (1996) Specific cleavage of α-fodrin during fas- and tumor necrosis factor-induced apoptosis is mediated by an interleukin-1β-converting enzyme/Ced-3 protease distinct from the poly(ADP-ribose) polymerase protease. J. Biol. Chem. 271, 31277–31282

51. Benson, D. L., Mandell, J. W., Shaw, G., and Banker, G. (1996) Compartmentation of alpha-internexin and neurofilament triplet proteins in cultured hippocampal neurons. J. Neurocytol. 25, 181–196

52. Tian, W., Czopka, T., and López-Schier, H. (2020) Systemic loss of Sarm1 protects Schwann cells from chemotoxicity by delaying axon degeneration. Commun. Biol. 3, 1–14

53. Mishra, B., Carson, R., Hume, R. I., and Collins, C. A. (2013) Sodium and potassium currents influence wallerian degeneration of injured Drosophila axons. J. Neurosci. 33, 18728–18739

54. Avery, M. A., Rooney, T. M., Pandya, J. D., Wishart, T. M., Gillingwater, T. H., Geddes, J. W., Sullivan, P. G., and Freeman, M. R. (2012) Wld S prevents axon degeneration through increased mitochondrial flux and enhanced mitochondrial Ca^2+^ buffering. Curr. Biol. 22, 596–600

55. Baccei, M. L., and Kocsis, J. D. (2000) Voltage-gated calcium currents in axotomized adult rat cutaneous afferent neurons. J. Neurophysiol. 83, 2227–2238

56. Fuchs, A., Rigaud, M., Sarantopoulos, C. D., Filip, P., and Hogan, Q. H. (2007) Contribution of calcium channel subtypes to the intracellular calcium signal in sensory neurons: The effect of injury. Anesthesiology. 107, 117–127

57. Sasaki, Y., Engber, T. M., Hughes, R. O., Figley, M. D., Wu, T., Bosanac, T., Devraj, R., Milbrandt, J., Krauss, R., and DiAntonio, A. (2020) cADPR is a gene dosage-sensitive biomarker of SARM1 activity in healthy, compromised, and degenerating axons. Exp. Neurol. 329, 113252

58. Li, Y., Pazyra-Murphy, M. F., Avizonis, D., Russo, M. T., Tang, S., Chen, C. Y., Hsueh, Y. P., Bergholz, J. S., Jiang, T., Zhao, J. J., Zhu, J., Ko, K. W., Milbrandt, J., Diantonio, A., and Segal, R. A. (2022) Sarm1 activation produces cADPR to increase intra-axonal Ca++ and promote axon degeneration in PIPN. J. Cell Biol. 10.1083/jcb.202106080

59. Essuman, K., Milbrandt, J., Dangl, J. L., and Nishimura, M. T. (2022) Shared TIR enzymatic functions regulate cell death and immunity across the tree of life. Science. 377, 1–21

60. Guse, A. H. (2015) Calcium mobilizing second messengers derived from NAD. Biochim. Biophys. Acta - Proteins Proteomics. 1854, 1132–1137

61. Lee, H. C., and Zhao, Y. J. (2019) Resolving the topological enigma in Ca2+ signaling by cyclic ADP-ribose and NAADP. J. Biol. Chem. 294, 19831–19843

62. Metwally, E., Zhao, G., and Zhang, Y. Q. (2021) The calcium-dependent protease calpain in neuronal remodeling and neurodegeneration. Trends Neurosci. 44, 741–752

63. Mingorance-Le Meur, A., and O’Connor, T. P. (2009) Neurite consolidation is an active process requiring constant repression of protrusive activity. EMBO J. 28, 248–260

64. Rueden, C. T., Schindelin, J., Hiner, M. C., DeZonia, B. E., Walter, A. E., Arena, E. T., and Eliceiri, K. W. (2017) ImageJ2: ImageJ for the next generation of scientific image data. BMC Bioinformatics. 18, 1–26

65. Chen, Z., Kwong, A. K. Y., Yang, Z., Zhang, L. L., Lee, H. C., and Zhang, L. L. (2011) Studies on the synthesis of nicotinamide nucleoside and nucleotide analogues and their inhibitions towards CD38 NADase. Heterocycles. 83, 2837–2850

66. Lenth, R. V. (2022) emmeans: Estimated marginal means, aka least-squares means

