## Supplemental Tables for "SARM1 is responsible for calpain-dependent dendrite degeneration in mouse hippocampal neurons"

**Table 1: Antibodies used**

| Target | Clone | Source | Identifier | RRID | Application | Concentration |
| --- | --- | --- | --- | --- | --- | --- |
| α-Internexin | Rb polyclonal | Novus | NB300-139 | AB_10001006 | ICC | 1:1000 |
| αII-Spectrin | 3D7 | Novus | NBP1-92689 | AB_11031596 | ICC | 1:500 |
| Axonal neurofilaments | SMI312 | BioLegend | 837904 | AB_2566782 | ICC | 1:2000 |
| MAP2 | Chk polyclonal | Abcam | ab5392 | AB_2138153 | ICC | 1:5000 |
| MAP2 | Rb polyclonal | Abcam | ab32454 | AB_776174 | ICC | 1:2000 |
| Nf-L | C28E10 |  | 2837 | AB_823575 | ICC | 1:1000 |
| Nf-M | Rb polyclonal | BioLegend | 841001 | AB_2565457 | ICC | 1:500 |
| SARM1 | W16079A | BioLegend | 696602 | AB_2687069 | ICC | 1:100 |
| Specific species | Various | Thermo Fisher Scientific | Secondary antibodies conjugated with Alexa Fluor dyes | Various | ICC | 1:1000 |
| α-Internexin | Rb polyclonal | CST | 77024 | na | WB | 1:2000 |
| αII-Spectrin | 3D7 | Novus | NBP1-92689 | AB_11031596 | WB | 1:2000 |
| Nf-L | Rb polyclonal | Novus | NB300-131 | AB_2251212 | WB | 1:20000 |
| Tuj1 | TUJ1 | BioLegend | 801202 | AB_10063408 | WB | 1:3000 |
| Specific species | Various | Li-cor biosciences | Secondary antibodies conjugated with IRDyes | Various | WB | 1:10000 |

**Table 2: Key Reagents**

| Acronym | Reagents | Source | Identifier |
| --- | --- | --- | --- |
| 1x PBS | PBS, pH7.4 | Thermo Fisher Scientific | 10010023 |
| 5-Fluoro-2’deoxyuridine | 5-Fluoro-2’deoxyuridine | Sigma-Aldrich | 856657 |
| B27 | B27 Supplement (50x), serum free | Thermo Fisher Scientific | 17504044 |
| Biocoat 96-well plates | BioCoat Poly-D-Lysine 96-well Black/Clear plates | Corning | 354640 |
| Biocoat 384-well plates | BioCoat Poly-D-Lysine 384-well Black/Clear flat bottom plates | Corning | 354663 |
| BSA | Bovine Serum Albumin | Sigma-Aldrich | A2153 |
| Calpain Inhibitor III | Calpain Inhibitor III / MDL 28170, CAS: 88191-84-8 | Sigma-Aldrich | 208722-25MG |
| Calpeptin | Calpeptin, CAS: 117591-20-5 | Tocris | 0448 |
| Cell Mask | CellMask Depp Red Plasma Membrane Stain | Thermo Fisher Scientific | C10046 |
| Cilnidipine | Cilnidipine, CAS: 132203-70-4 | Tocris | 2629 |
| CZ-48 | CZ-48, CAS: 1374663-29-2 | Synthesized. See Methods | CAS: 1374663-29-2 |
| DAPI | DAPI solution (1mg/ml) | Thermo Fisher Scientific | 62248 |
| DNAseI | Deoxyribonuclease I from bovine pancreas | Sigma-Aldrich | D5025-150KU |
| EGTA | EGTA 1M, pH8.0, Sterile | BioWorld | 40120128-1 |
| EST | EST, CAS: 88321-09-9 | Sigma-Aldrich | 330005-1MG |
| Fluo4 | Fluo-4, AM | Thermo Fisher Scientific | F14201 |
| FK866 | FK 866 hydrochloride, CAS: 2727965-45-7 | Tocris | 4808 |
| Glucose | 1M Glucose | Boston BioProducts | BM-675 |
| Glutamax | GlutaMAX^TM^ Supplement | Thermo Fisher Scientific | 35050061 |
| HI-FBS | Heat-inactivated fatal bovine serum | VWR | 97068-085 |
| Hibernate E | Hibernate E - Calcium -Magnesium | BrainBits | HECAMG500 |
| L15 | Leibovitz’s L-15 medium | Thermo Fisher Scientific | 11415064 |
| Laminin | Natural Mouse Laminin | Thermo Fisher Scientific | 23017015 |
| Leupeptin | Leupeptin, CAS: 103476-89-7 | Enzo Life Sciences | ALX-260-009-M005 |
| ML218 | ML218 hydrochloride, CAS: 2319922-08-0 | Tocris | 4507 |
| NAD/NADH-Glo assay | NAD/NADH-Glo assay | Promega | G9072 |
| NbActiv4 | NbActiv4 (for primary neuron cultures) | BrainBits | Nb4-500 |
| Neurobasal medium | Neurobasal Medium | Thermo Fisher Scientific | 21103049 |
| NGF | Nerve Growth Factor, NGF 2.5S Murine | Promega | G514A |
| Nifedipine | Nifedipine, CAS: 21829-25-4 | Tocris | 1075 |
| NuPAGE LDS sample buffer | NuPAGE LDS Sample Buffer (4x) | Thermo Fisher Scientific | NP0007 |
| NuPAGE MOPS SDS running buffer | NuPAGE MOPS SDS running buffer | Thermo Fisher Scientific | MP000102 |
| NuPAGE Novex 4-12% Bis-Tris Midi protein gels | NuPAGE Novex 4-12% Bis-Tris Midi protein gels | Thermo Fisher Scientific | WG1403 |
| NuPAGE Sample Reducing Agent | NuPAGE Sample Reducing Agent (10x) | Thermo Fisher Scientific | NP0009 |
| Papain | PDS Kit, Papain vial | Worthington | LK003178 |
| Paraformaldehyde (PFA) | 32% PFA | Electron Microscopy Sciences | 15714-S |
| PC6 | PC6 | Synthesized. See Methods | (Li et al., 2021) |
| Penicillin-Streptomycin | Penicillin-Streptomycin (10000U/ml) | Thermo Fisher Scientific | 15140122 |
| PD150606 | PD150606, CAS: 426821-41-2 | Sigma-Aldrich | 513022-5MG |
| Q-VD-OPh | Q-VD-OPh hydrate | Sigma-Aldrich | SML0063-1MG |
| Sucrose | Sucrose | Sigma-Aldrich | S0389 |
| TBST | Tris-buffered saline with 0.5% Tween20 | Teknova | T9501 |
| Trans-Blot Turbo Transfer System | Trans-Blot Turbo Transfer System, 0.2µm Nitrocellulose | Bio-Rad | 1704270 |
| SKF96365 | SKF 96365 hydrochloride, CAS: 130495-35-1 | Tocris | 1147 |
| Triton X100 | Triton X-100 | Thermo Fisher Scientific | BP151-500 |
| Vacor | Pyrinuron, CAS: 53558-25-1 | Chem Service | N-13738 |
